# Astrocyte growth during morphogenesis is driven by the Tre1/S1pr1 phospholipid-binding G protein-coupled receptor

**DOI:** 10.1101/2022.09.15.508188

**Authors:** Jiakun Chen, Tobias Stork, Yunsik Kang, Amy Sheehan, Cameron Paton, Kelly R. Monk, Marc R. Freeman

## Abstract

Astrocytes play crucial roles in regulating neural circuit function by forming a dense network of synapse-associated membrane specializations, but signaling pathways regulating astrocyte morphogenesis remain poorly defined. Here we show the *Drosophila* lipid-binding G protein-coupled receptor (GPCR) Tre1, likely acting through Rac1, is required for astrocytes to elaborate their complex morphology *in vivo*. The lipid phosphate phosphatases Wunen/Wunen2, which process phospholipid ligands, also regulate astrocyte morphology, and, via Tre1, mediate astrocyte-astrocyte competition for growth promoting lipids. Loss of *s1pr1*, the functional analog of *Tre1* in zebrafish disrupts astrocyte process elaboration. Live-imaging and pharmacology demonstrate that S1pr1 balances proper astrocyte process extension/retraction dynamics during morphogenesis, and that S1pr1 signaling is required throughout astrocyte development. Tre1 and S1pr1 are thus potent evolutionarily conserved regulators of astrocyte growth and elaboration of morphological complexity.

- The GPCR Tre1 and LPPs Wun/Wun2 promote astrocyte process outgrowth in *Drosophila*
- Astrocytes compete for a growth­promoting phospholipid in the CNS
- Wun/Wun2 act locally to regulate process outgrowth through Tre1
- Vertebrate S1pr1 regulates astrocyte growth early, through modulation of process dynamics

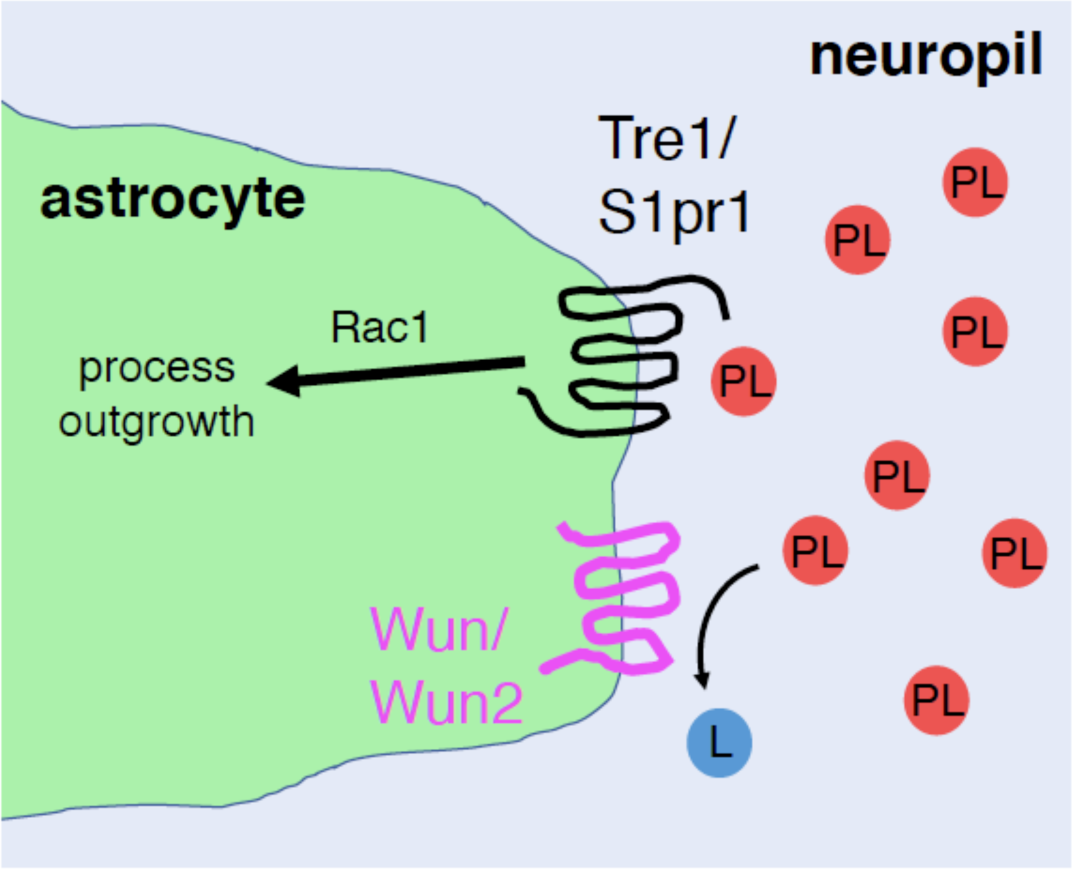

## INTRODUCTION

Astrocytes are essential for central nervous system (CNS) development, function and homeostasis across species. Astrocytes actively participate in synapse formation, pruning, and plasticity during circuit development (Allen et al., 2012; Christopherson et al., 2005; Chung et al., 2013; Lee et al., 2021). In myriad ways, astrocytes support neural circuit function by maintaining an optimized environment for neuronal activity (Allen and Barres, 2009; Freeman and Doherty, 2006; Khakh and Sofroniew, 2015). Increasing evidence also argues that astrocytes can modulate neural circuit activity and behavior, in some cases by being directly integrated into circuit signaling events (Bazargani and Attwell, 2016; Nagai et al., 2021). A single mammalian astrocyte can interact with thousands of synapses and physiologically bridge them with other CNS cell types. Most astrocyte functions are believed to depend on their profuse membrane specializations and morphological complexity, which act to facilitation of widespread signaling across different synapses, the vasculature, and other neighboring CNS cells. Dysregulated astrocyte morphology is a hallmark of many neuroinflammatory and neurological diseases (Burda and Sofroniew, 2014; Han et al., 2021; Molofsky et al., 2012), but we understand little about the mechanistic basis of astrocyte morphogenesis *in vivo*.

During development astrocytes acquire their elaborate morphology through a stepwise process of vigorous cellular process outgrowth to cover the CNS synaptic space, tiling with adjacent astrocytes, and extension of fine membrane leaflets adjacent to synapses (Allen and Barres, 2009). The Heartless/FGF receptor signaling in *Drosophila* acts mainly in a permissive fashion to enable astrocyte process infiltration of the synaptic neuropil, but when FGF ligands are ectopically expressed, astrocyte membranes can be redirected to inappropriate regions of the CNS (Stork et al., 2014). Astrocyte tiling in mammals occurs after initial overlap of immature astrocyte territories, followed by the establishment of more precise non-overlapping domains (Bushong et al., 2004). The boundaries of astrocyte domains are believed to develop through repulsive or competitive interactions with neighboring astrocytes. This notion is supported by the minimal overlap observed between astrocytes in mammals (Bushong et al., 2002), zebrafish (Chen et al., 2020), and *Drosophila* (Stork *et al*., 2014), and the observation that ablation of a subset of *Drosophila* astrocytes during development leads to the expansion of remaining astrocyte processes into unoccupied regions of the neuropil (Stork *et al*., 2014). In mammals, loss of the cell adhesion molecule HepaCAM reduces astrocyte domain volume, morphological complexity, and astrocyte gap junction coupling (Baldwin et al., 2021b), arguing for an important role for HepaCAM in astrocyte morphogenesis. Similarly, the dense network of fine astrocyte leaflets is reduced in animals lacking Neurexin/Neuroligin (Stogsdill et al., 2017), which likely serves to coordinate astrocyte-synapse contact sites and synaptogenesis. Astrocyte morphogenesis occurs in parallel with the wiring of neural circuits, and both are intertwined, as astrocyte depletion during development, or the elimination of astrocyte-expressed molecules results in profound abnormalities in neural circuits, including reduced synapse numbers (Ackerman et al., 2021; Allen *et al*., 2012; Christopherson *et al*., 2005; Delaney et al., 1996; Eroglu et al., 2009; Kucukdereli et al., 2011; Muthukumar et al., 2014; Tsai et al., 2012).

The *Drosophila* gene *Tre1* (*Trapped in endoderm 1*) encodes a lipid-binding G protein-coupled receptor (GPCR) that promotes the survival and transepithelial migration of germ cells. Maternal *Tre1* RNA is localized to germ cells where it functions cell-autonomously (Kunwar et al., 2003). Wunen (Wun) and Wunen2 (Wun2) are lipid phosphate phosphatases (LPPs), membrane-embedded enzymes that regulate levels of extracellular bioactive lipids such as sphingosine-1-phosphate (S1P) and lysophosphatidic acid (Morris et al., 2013). Tre1 is thought to be activated by a phospholipid ligand, which is processed by Wun and/or Wun2 based on the observations that *wun/wun2* mutants partially phenocopy *Tre1* mutants and genetic interaction studies (LeBlanc and Lehmann, 2017; Zhang et al., 1997). Tre1 belongs to a family of rhodopsin-like GPCRs, which contain two highly conserved domains (NRY and NPxxY motifs), and each domain mediates distinct downstream signaling events (LeBlanc and Lehmann, 2017). While the phospholipid ligand acting on Tre1 in *Drosophila* germ cell migration is not known, S1P and lipid-binding GPCRs like the S1P receptor 1 (S1pr1) appear to have conserved roles in germ cell migration (Kassmer et al., 2015; Richardson and Lehmann, 2010).

In this study, we show lack of Tre1 or the LPPs Wun/Wun2 in astrocytes dramatically reduces the morphological complexity of fine astrocyte processes in *Drosophila*. Tre1 signals through the small GTPase Rac1, modulating astrocyte process dynamics during outgrowth and organization of the astrocyte cytoskeleton. Wun/Wun2 act in astrocytes to promote astrocyte process outgrowth in a Tre1-dependent fashion. Wun/Wun2 can be repulsive to astrocyte processes when expressed in neurons and can disrupt astrocyte-astrocyte competition when differentially expressed in adjacent astrocytes, arguing for a role for this pathway in establishing individual astrocyte morphological domains. Genetic or pharmacological blockade of S1pr1 signaling in zebrafish similarly perturbs astrocyte growth, and live-imaging of the earliest stages of astrocyte development reveals a key role for S1pr1 in astrocyte process extension/retraction dynamics. Our work thus identifies Tre1/S1pr1 and extracellular phospholipids as key regulators of astrocyte morphological development from flies to vertebrates.

## RESULTS

### *Drosophila Tre1* mutants have smaller astrocytes with reduced morphological complexity

To identify genes enriched in astrocytes, we analyzed existing gene expression databases in *Drosophila* (Brunet Avalos et al., 2019; Davie et al., 2018; Huang et al., 2015; Ng et al., 2016). We found that *Tre1* expression is considerably higher in astrocytes than that detected in total brain cells, and in an unrelated screen for genes required for larval astrocyte engulfment function, we found that expression of a *Tre1^RNAi^* construct in glial cells led to gross defects in astrocyte morphology. To more rigorously test the role of Tre1 in astrocyte morphogenesis, we examined astrocyte morphology in various *Tre1* mutants, in particular the *Tre1^attP^* null mutant in which the genomic coding region of *Tre1* is replaced with an *attP* docking site (Deng et al., 2019; see also Figures S1A and S1B). We labeled adult astrocyte membranes with anti-Gat antibody and presynaptic active zones in the neuropil with anti-Bruchpilot (Brp). In *Tre1^attP^* mutants, we found that astrocyte membrane coverage of the antennal lobe (AL) neuropil was dramatically decreased. Astrocyte processes in *Tre1^attP^* mutants were thicker than in controls, and did not fully elaborate fine processes to infiltrate olfactory glomeruli (Figure 1A). We found similar phenotypes in *Tre1^sctt^* mutants (Figure S1C), which results from a mis-splicing of the *Tre1* transcript in the coding region (Kamps et al., 2010). However, we did not observe obvious phenotypes in *Tre1*^D^*^EP5^* mutants (Figure S1C), in which the non-coding first exon region of *Tre1* is deleted (Kunwar *et al*., 2003). The lack of phenotypic effects of the *Tre1*^D^*^EP5^* mutant is likely due to the use of an alternative *Tre1* transcriptional start in astrocytes that is not affected by the *Tre1*^D^*^EP5^* mutation (Figure S1A).

**Figure 1.**
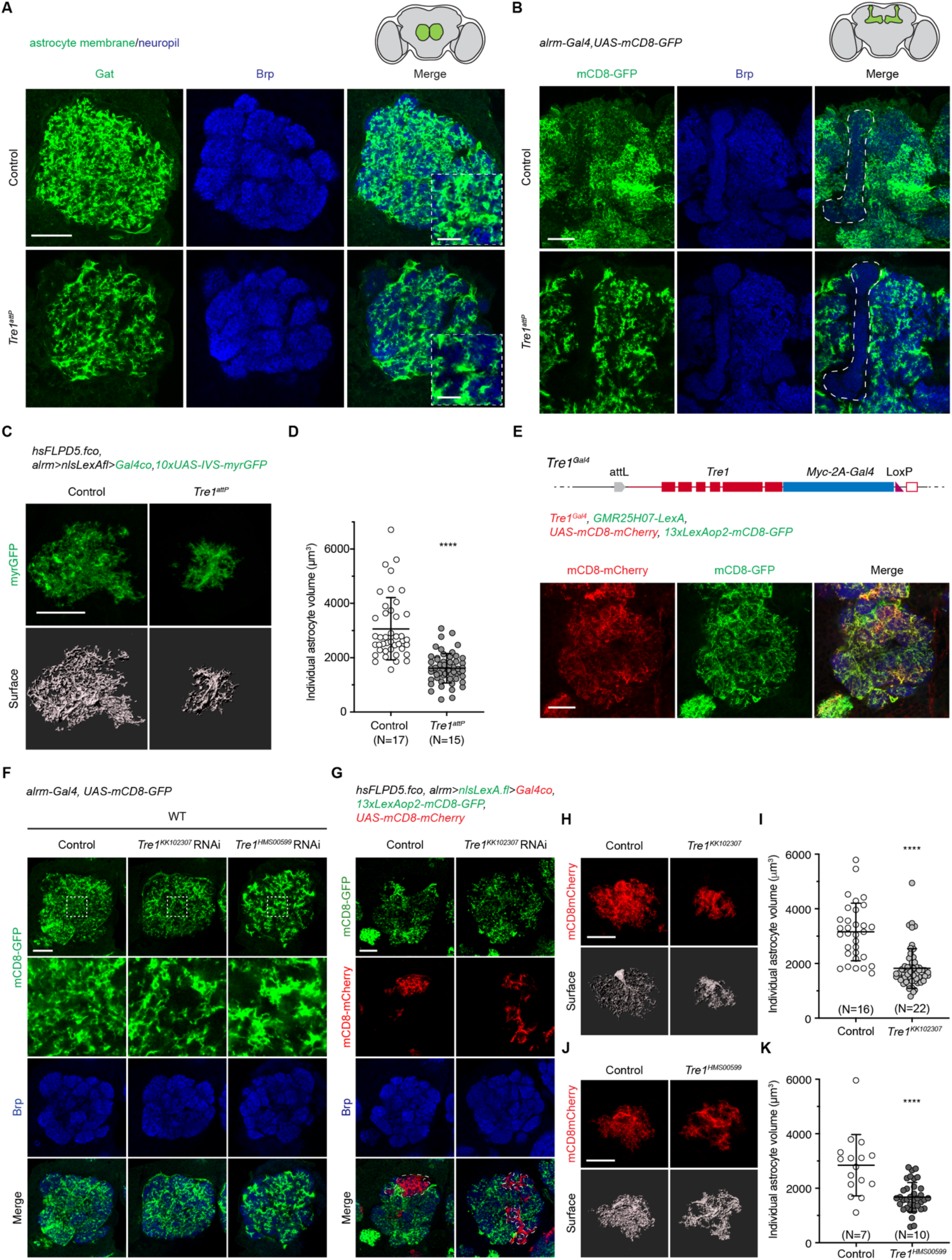
*Drosophila* Tre1 regulates astrocyte fine process infiltration autonomously in astrocytes. (A) Astrocytes labeled with membrane marker α-Gat antibody (green) and synaptic neuropil with α-Brp (blue) in the adult *Drosophila* brain antennal lobe (AL) in control (N=5) and *Tre1^attP^* mutants (N=8). Insets show enlarged views to highlight the astrocyte membrane infiltration differences observed in control and *Tre1^attP^* mutants. N, number of animals. Scale bars, 20 µm and 5 µm (insets). (B) *alrm-Gal4 UAS-mCD8-GFP*-labeled astrocytes (green) and the neuropil (blue) in control (N=5) and *Tre1^attP^* mutant (N=4) protocerebrum. Dashed lines mark the mushroom body. (C) Images of single-cell astrocytes labeled with membrane marker myrGFP (green) using the FLP-out system and the representative IMARIS 3D-rendering surface (grey) in control and *Tre1^attP^* mutants. Scale bar, 20 µm. (D) Quantification of individual astrocyte volumes in control (N=17) and *Tre1^attP^* mutants (N=15). Data points represent single astrocytes. ****, p<0.0001; unpaired t test. Error bars, mean values ± S.D. (E) *Tre1^Gal4^*-expressing mCD8-mCherry cell membrane (red) with astrocyte-specific *GMR25H07-LexA 13xLexAop2-mCD8-GFP*-labeled astrocyte membrane (green) in the ALs (N=7), and the synaptic neuropil is labeled by α-Brp (blue). Scale bar, 20 µm. (F) Images of AL astrocyte membrane surface labeled with alrm-Gal4 UAS-mCD8-GFP (green) and neuropil by α-Brp (blue) in control (N=9), *Tre1^KK102307^ RNAi* (N=10), and *Tre1^HMS00599^ RNAi* (N=10). Scale bar, 20 µm. (G) Images of FLP-out approach generated mosaic astrocyte clones labeled by mCD8-mCherry (red) in control or *Tre1^KK102307^ RNAi* in adjacent to neighboring WT mCD8-GFP-expressing astrocytes (green). Dashed lines mark the boundaries of mCD8-mCherry-labeled astrocytes at a single-Z plane. Scale bar, 20 µm. (H) Images of single-cell astrocytes labeled with mCD8-mCherry (red) and the representative IMARIS 3D-rendering surface (grey) in control and *Tre1^KK102307^ RNAi*. Scale bar, 20 µm. (I) Quantification of individual astrocyte volumes in control (N=16) and *Tre1^KK102307^ RNAi* (N=22). Data points represent single astrocytes. ****, p<0.0001; unpaired t test. Error bars, mean values ± S.D. (J) Images of single-cell astrocytes labeled with mCD8-mCherry (red) and the representative IMARIS 3D-rendering surface (grey) in control and *Tre1^HMS00599^ RNAi*. Scale bar, 20 µm. (K) Quantification of individual astrocyte volumes in control (N=7) and *Tre1^HMS00599^ RNAi* (N=10). Data points represent single astrocytes. ****, p<0.0001; unpaired t test. Error bars, mean values ± S.D. See also Figure S1.

We next analyzed astrocyte morphology using the astrocyte-specific driver *alrm-Gal4* to drive *UAS-mCD8-GFP* to label astrocyte membranes. We observed reduced astrocyte infiltration across the entire adult CNS in *Tre1^attP^* mutants. In the mushroom body, where wild-type astrocytes show a slightly lower density of processes compared to other regions of the neuropil (Kremer et al., 2017), *Tre1^attP^* mutants exhibit a near complete absence of astrocyte processes (Figure 1B). Defects in astrocyte infiltration were present beginning during larval stages (Figure S1D) and persisted through adulthood, indicating that Tre1 is required throughout development (Figure S1D). To determine precisely how individual astrocytes were disrupted in the *Tre1^attP^* mutants, we used a FLP-out strategy to generate single-cell astrocyte clones labeled with myristoylated-GFP. We found that compared to controls, *Tre1^attP^* mutant astrocytes occupied a smaller territory in the synaptic neuropil, and the total cell volume was significantly decreased in comparison with WT control astrocytes (Figures 1C and 1D). Taken together, these data indicate that *Tre1* is required for normal astrocyte morphogenesis.

### Tre1 acts cell autonomously in astrocytes to regulate membrane growth

To explore the endogenous expression pattern of *Tre1* in the adult brain, we took advantage of the *Tre1^attP^* mutation, where an *attP* docking site is inserted at the *Tre1* genomic locus, to generate a *Tre1^Gal4^* knock-in allele, in which *Gal4* is co-expressed together with endogenous *Tre1* by a self-cleaving T2A peptide sequence (Figure S1E). *Tre1^Gal4^*-driven mCD8-mCherry distribution overlapped with Gat-labeled astrocyte membrane in the ALs, and most of the lam-GFP-labeled nuclei were surrounded by anti-Gat staining, indicating they were astrocyte cell bodies (Figure S1F). We next compared *Tre1^Gal4^>mCD8-mCherry* expression with *GMR25H07-LexA>mCD8-GFP*, where astrocytes express the membrane marker mCD8-GFP under *LexA/LexAop* control. We observed that *Tre1^Gal4^*-controlled mCD8-mCherry labeling overlapped with mCD8-GFP-labeled astrocyte membranes throughout the adult brain (Figures 1E and S1G). These data, combined with transcriptomic data, show that *Tre1* expression is highly enriched in *Drosophila* astrocytes.

To determine the cell-autonomy of Tre1 function, we first used the *alrm-Gal4* driver to express two independent, non-overlapping *UAS-Tre1* RNAi constructs. By knocking down *Tre1* in astrocytes, the coverage of AL neuropil by astrocyte processes was reduced in both *Tre1* RNAi conditions in contrast to the controls (Figure 1F). We next generated single-cell RNAi clones using the FLP-out system, where WT astrocytes labeled with mCD8-GFP were adjacent to mCD8-mCherry-labeled clones that were either controls (no RNAi) or expressed *Tre1^RNAi^*. In control experiments, both the mCD8-GFP- and the mCD8-mCherry-labeled WT astrocytes exhibited dense infiltration of the neuropil, and individual astrocytes (mCD8-mCherry^+^) occupied unique territories that did not overlap with neighboring cells (Figure 1G). In contrast, when we generated astrocytes expressing *Tre1 RNAi* constructs, the mCD8-mCherry^+^ clones displayed decreased density of processes, with gaps in-between main branches (Figure 1G). We often observed that these gaps were invaded by neighboring mCD8-GFP-labeled WT astrocytes (Figure 1G). Moreover, individual *Tre1^RNAi^* astrocytes showed a strongly reduced total volume in comparison with controls (Figures 1H-K). We conclude that *Tre1* is required in astrocytes for normal growth of astrocyte volume and fine leaflets, and that WT astrocytes can compensate for gaps in *Tre1^RNAi^* astrocytes to ensure full coverage of the neuropil.

### Tre1 drives astrocyte growth through its NPIIY motif

In *Drosophila* germ cells, the two conserved NRY and NPIIY motifs of Tre1 are required to guide proper germ cell migration (LeBlanc and Lehmann, 2017). We generated *UAS-Tre1* rescue constructs with either wild-type *Tre1*, single mutations specific to each domain, or double mutations (*Tre1^NAY^*, *Tre1^AAIIY^*, and *Tre1^NAY,AAIIY^,* (LeBlanc and Lehmann, 2017))(Figure 2A). We assayed for rescue of *Tre1^attP^* mutant phenotypes by driving their expression in astrocytes. We found that both wild-type *Tre1* and *Tre1^NAY^* transgenes rescued astrocytic coverage of the neuropil in the *Tre1^attP^* mutants at levels comparable to that observed in control astrocytes (Figure 2B). In contrast, neither *Tre1^AAIIY^* nor *Tre1^NAY,AAIIY^* transgenes could suppress the infiltration defects in the *Tre1^attP^* mutant backgrounds (Figure 2B).

**Figure 2.**
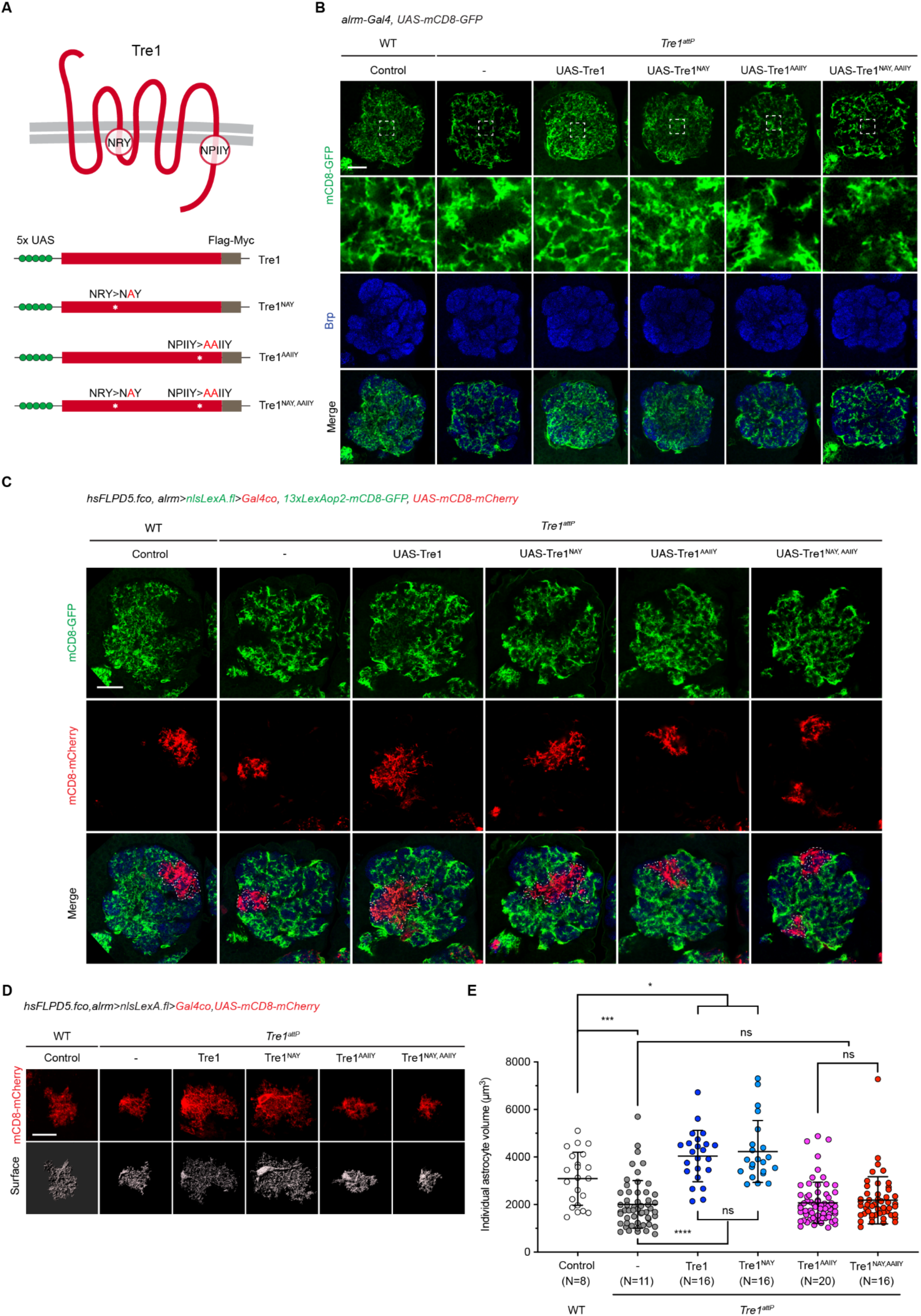
Tre1 controls astrocyte morphology via the NPIIY motif but not the NRY motif. (A) Schematics of Tre1 GPCR with the conserved NRY and NPIIY motifs and the *UAS*-driven transgenic constructs. (B) Images of AL astrocytes labeled by *alrm-Gal4 UAS-mCD8-GFP* (green) and the neuropil by α-Brp (blue) in WT control (N=6) and in the *Tre1^attP^* mutant backgrounds either alone (N=11) or that co-express *UAS-Tre1* (N=8), *UAS-Tre1^NAY^* (N=8), *UAS-Tre1^AAIIY^* (N=11), *UAS-Tre1^NAY,AAIIY^* (N=10). Insets show enlarged views to highlight the astrocyte membrane infiltration differences observed in different experimental conditions. Scale bars, 20 µm and 5 µm (insets). (C) Images of astrocytes labeled by mCD8-GFP (green) with FLP-out clones expressing mCD8-mCherry (red) in the contexts of WT control or *Tre1^attP^* mutants that also express various forms of *Tre1* transgenic constructs. Dashed lines mark the boundaries of mCD8-mCherry-labeled astrocytes at a single-Z plane. Scale bar, 20 µm. (D) Images of single-cell astrocytes labeled with mCD8-mCherry (red) and the representative IMARIS 3D-rendering surface (grey) in WT control and in the *Tre1^attP^* mutant backgrounds that express various forms of *Tre1* transgenic constructs. Scale bar, 20 µm. (E) Quantification of individual astrocyte volumes in the corresponding experimental conditions as shown in (D). N, number of animals. WT control, N=8; *Tre1^attP^*, N=11; *Tre1^attP^* + *Tre1*, N=16; *Tre1^attP^* + *Tre1^NAY^*, N=16; *Tre1^attP^* + *Tre1^AAIIY^*, N=20; *Tre1^attP^* + *Tre1^NAY,AAIIY^*, N=16. Data points represent single astrocytes. ns, not significant; *, p<0.05; ***, p<0.001; ****, p<0.0001; one-way ANOVA with multiple comparisons. Error bars, mean values ± S.D. See also Figure S2.

Using the FLP-out strategy, we analyzed single-cell clones in *Tre1^attP^* mutant backgrounds. We found that both wild-type *Tre1*- and *Tre1^NAY^*-expressing astrocyte clones exhibited a WT-like dense infiltration of their processes into the neuropil (Figure 2C), and individual cell volumes were rescued compared to *Tre1* mutant clones (Figures 2D and 2E). Interestingly, we also found that individual cell volumes in both wild-type *Tre1*- and *Tre1^NAY^*-expressing astrocyte clones in *Tre1^attP^* mutants were significantly larger than WT control astrocyte clones (Figures 2D and 2E), suggesting that Tre1 overexpression can increase astrocyte clone size, which is perhaps enhanced by the presence of neighboring *Tre1*-deficient astrocytes. However, expression of *Tre1^AAIIY^* or *Tre1^NAY,AAIIY^* transgenes failed to rescue *Tre1^attP^* mutant phenotypes (Figures 2C-E). In addition, overexpressing these constructs in astrocytes did not alter astrocyte growth in a WT background, suggesting none of these transgenes dominantly interfere with native Tre1 function (Figures S2A and S2B). Compared to wild-type Tre1 or Tre1^NAY^, Tre1 proteins with a mutated NPIIY motif (Tre1^AAIIY^ and Tre1^NAY,AAIIY^) were less distributed into astrocyte processes and showed accumulation in cell bodies, suggesting a role for the NPIIY motif in Tre1 trafficking or turnover (Figure S2C). These results indicate that the Tre1 NPIIY motif, but not the NRY motif, is required for astrocyte morphogenesis, and suggest that the downstream signaling cascades activated by Tre1 in astrocytes are different from those in germ cells where both the NRY and NPIIY motifs drive important signaling events (LeBlanc and Lehmann, 2017).

Rac1 is essential for nervous system development and well-known for its role in actin cytoskeleton regulation (Luo et al., 1994; Ng et al., 2002), and *in vitro* studies demonstrate Rac1 can contribute to astrocyte morphological changes (Racchetti et al., 2012). We used three different *Rac1* transgenic lines to explore potential interactions with *Tre1*: *UAS-Rac1.V12* (a constitutively active form of Rac1), *UAS-Rac.N17* (a dominant-negative form of Rac1), and *UAS-Rac1.W* (a wild-type form of Rac1). Expressing constitutively active *Rac1.V12* in astrocytes led to cell death, therefore, we focused our analyses on the other two forms of Rac1. Using the FLP-out system in a WT background, when Rac1.N17 or Rac1.W was misexpressed in single-cell clones labeled with mCD8-mCherry, astrocyte morphology displayed reduced fine process elaboration (Figure 3A), and individual astrocytes showed decreased cell volumes (Figures 3B and 3C). These data suggest that both loss- and gain-of-function in Rac1 signaling are disruptive to astrocyte morphogenesis, and that Rac1 activity needs to be finely tuned for normal astrocyte growth. To test whether *Tre1* genetically interacts with *Rac1*, we next manipulated Rac1 activity in the *Tre1^attP^* mutant backgrounds. Both Rac1.N17 and Rac1.W expression in astrocytes resulted in an enhancement of defects in astrocyte phenotypes compared to *Tre1^attP^* mutants alone (Figure 3D). Individual mCD8-mCherry^+^ *Tre1* mutant astrocytes co-expressing Rac1.N17 or Rac1.W extended almost no fine membrane processes and their astrocyte volumes were highly reduced (Figures 3E-G). Together, these observations support the notion that Tre1 functions to regulate Rac1 activity during astrocyte morphogenesis.

**Figure 3.**
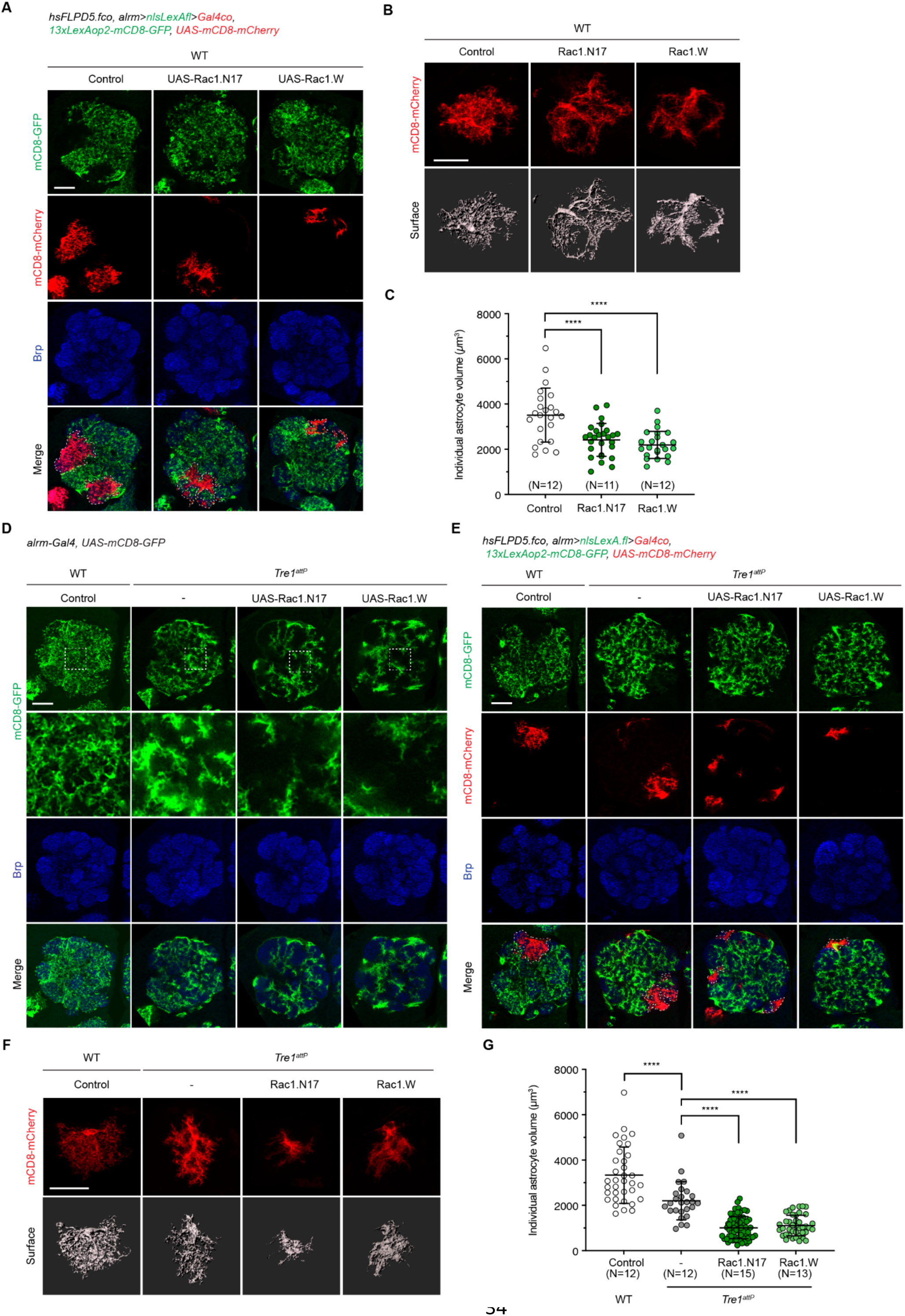
Tre1 balances Rac1 activity to govern proper astrocyte growth. (A) Images of astrocytes labeled by mCD8-GFP (green) with FLP-out clones expressing mCD8-mCherry (red) in the WT backgrounds that also express *UAS-Rac1.N17* or *UAS-Rac1.W*, and Brp labels the neuropil (blue). Dashed lines mark the boundaries of mCD8-mCherry-labeled astrocytes at a single-Z plane. Scale bar, 20 µm. (B) Images of single-cell astrocytes labeled with mCD8-mCherry (red) and the representative IMARIS 3D-rendering surface (grey) in WT backgrounds that express *Rac1.N17* or *Rac1.W*. Scale bar, 20 µm. (C) Quantification of individual astrocyte volumes in (B). N, number of animals. WT control, N=12; *Rac1.N17*, N=11; *Rac1.W*, N=12. Data points represent each single astrocyte. ****, p<0.0001; one-way ANOVA with Multiple comparisons. Error bars, mean values ± S.D. (D) Images of AL astrocyte membrane labeled with *alrm-Gal4 UAS-mCD8-GFP* (green) and neuropil with α-Brp (blue) in WT control (N=8), and in the *Tre1^attP^* mutant backgrounds that either alone (N=8) or co-express *UAS-Rac1.N17* (N=7), *UAS-Rac1.W* (N=8). N, number of animals. Scale bar, 20 µm. (E) Images of astrocytes labeled by mCD8-GFP (green) with FLP-out clones expressing mCD8-mCherry (red) in the contexts of WT control or *Tre1^attP^* mutants that also express *UAS-Rac1.N17* or *UAS-Rac1.W*. Dashed lines mark the boundaries of mCD8-mCherry-labeled astrocytes at a single-Z plane. Scale bar, 20 µm. (F) Images of single-cell astrocytes labeled with mCD8-mCherry (red) and the representative IMARIS 3D-rendering surface (grey) in WT control and in the *Tre1^attP^* mutant backgrounds that express *Rac1.N17* or *Rac1.W*. Scale bar, 20 µm. (G) Quantification of individual astrocyte volumes in (F) N, number of animals. WT control, N=12; *Tre1^attP^*, N=12; *Tre1^attP^* + *Rac1.N17*, N=15; *Tre1^attP^* + *Rac1.W*, N=13. Data points represent single astrocytes. ****, p<0.0001; one-way ANOVA with multiple comparisons. Error bars, mean values ± S.D.

### Tre1 and Rac1 regulate the astrocyte actin cytoskeleton

We next analyzed astrocyte cytoskeletal organization with cytoskeletal markers Lifeact-GFP.W and chRFP-Tub. We found the actin-rich fine cellular processes and the tubulin-rich major branches were both disrupted in the absence of *Tre1* (Figures S3A and S3B). Using Lifeact-GFP.W, we analyzed the morphological branching pattern of astrocytes at single-cell resolution using Imaris software (Figures 4A and S3C-D; Video S1). While the branch level of individual astrocytes in controls and *Tre1^attP^* mutants was comparable (Figures 4B and S3E), the branch points and total filament length were significantly reduced in the *Tre1^attP^* mutants (Figures 4C-D and S3F-G), and *Tre1^attP^* mutant astrocytes had an increased percentage of larger filament diameters (Figures 4E and S3H), consistent with the thicker processes we observed (Figure 1A). We found similar results when *Tre1^RNAi^* was driven in astrocytes (Figures S3I-M), and *Tre1^attP^* mutant morphological defects could be rescued by re-expression of wild-type *Tre1* in astrocytes, but not by *Tre1^NAY,AAIIY^* (Figures 4A-E). Tre1 thus plays a cell-autonomous role in regulating cytoskeletal organization in astrocyte fine processes, and loss of *Tre1* leads to a simplified astrocyte morphology with diminished branching and thicker processes.

**Figure 4.**
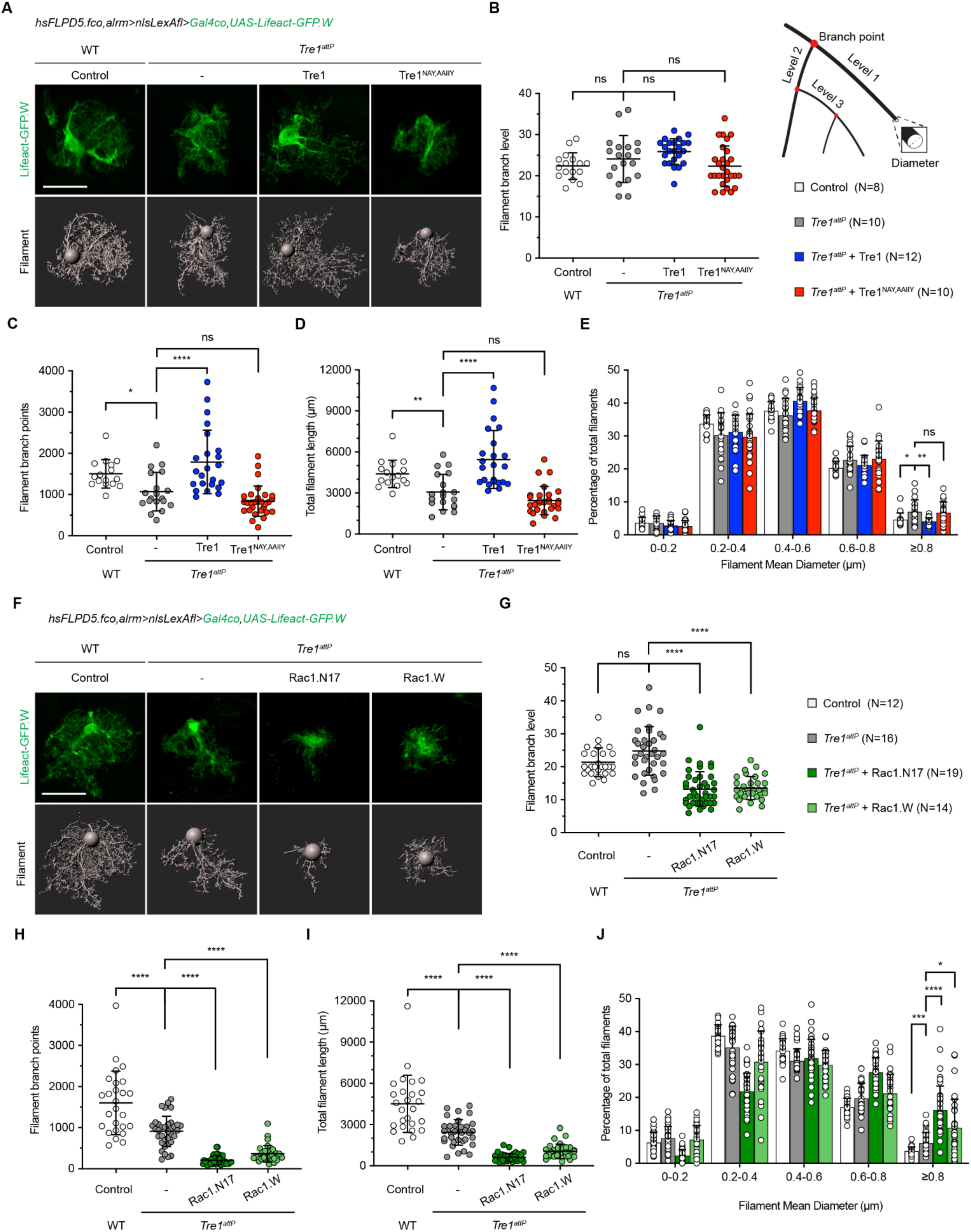
Tre1 regulates the actin cytoskeleton during astrocyte morphogenesis. (A) Images of single-cell astrocytes labeled with actin marker Lifeact-GFP.W (green) and the representative IMARIS 3D-rendering filament structures (grey) in WT control and in the *Tre1^attP^* mutant backgrounds either alone or that co-express *Tre1* and *Tre1^NAY,AAIIY^* constructs. Scale bar, 20 µm. (B-E) Quantification of single-cell astrocyte actin cytoskeletal branch level (B), branch points (C), total length (D), and distribution of mean diameter (E) in WT control (N=8), *Tre1^attP^* (N=10), *Tre1^attP^* + *Tre1* (N=12), and *Tre1^attP^* + *Tre1^NAY,AAIIY^* (N=10). N, number of animals. Data points represent single astrocytes. ns, not significant; *, p<0.05; **, p<0.01; ****, p<0.0001; one-way ANOVA with multiple comparisons. Error bars, mean values ± S.D. (F) Images of single-cell astrocytes labeled with Lifeact-GFP.W (green) and the representative IMARIS 3D-rendering filament structures (grey) in WT control and in the *Tre1^attP^* mutant backgrounds that express *Rac1.N17* or *Rac1.W*. Scale bar, 20 µm. (G-J) Quantification of single-cell astrocyte cytoskeletal branch level (G), branch points (H), total length (I), and distribution of mean diameter (J) in WT control (N=12), *Tre1^attP^* (N=16), *Tre1^attP^* + *Rac1.N17* (N=19), and *Tre1^attP^* + *Rac1.W* (N=14). N, number of animals. Data points represent single astrocytes. ns, not significant; *, p<0.05; ***, p<0.001; ****, p<0.0001; one-way ANOVA with Multiple comparisons. Error bars, mean values ± S.D. See also Figure S3, S4, and Video S1.

We next assayed the actin cytoskeletal phenotypes in Rac1 misexpression experiments. In control backgrounds, we found that inhibiting Rac1 activity in astrocytes with Rac1.N17 affected all aspects of cytoskeletal complexity, including decreases in branch level, branch points, and total filament length (Figures S4A-D) and an increase in larger filament diameter population (Figure S4E). However, astrocytic expression of Rac1.W did not influence branch level or the filament diameter distribution, but significantly decreased branch points and total filament length (Figures S4A-E). Moreover, by manipulating Rac1 activity in *Tre1^attP^* mutants, we observed a strong enhancement of actin cytoskeleton defects (actin branch level, branch points, total filament length, and distribution of filament diameter) by both Rac1.N17 and Rac1.W (Figures 4F-J). Together, these results support the notion that Tre1 regulates astrocyte cytoskeletal organization via Rac1.

### The lipid-binding GPCR S1pr1 regulates vertebrate astrocyte morphology

GPCRs and extracellular phospholipids control germ cell migration across species (Kassmer *et al*., 2015; Richardson and Lehmann, 2010). We sought to determine whether a lipid-binding GPCR plays a conserved role in vertebrate astrocyte morphogenesis similar to *Drosophila* Tre1. Zebrafish astrocytes are molecularly and functionally similar to their counterparts in *Drosophila* and mammals, and allow for direct visualization of astrocyte morphogenesis throughout development (Chen *et al*., 2020). We first assayed for phenotypes after loss of *Gpr84,* the most closely related vertebrate ortholog of *Drosophila Tre1* by sequence comparison (Figure S5A). We generated a mutation that resulted in 372 base-pair deletion of zebrafish *gpr84* (*gpr84^vo87^*) using CRISPR/Cas9-mediated genome editing (Jao et al., 2013; Jinek et al., 2012) (Figure S5B). The *gpr84^vo87^* mutation causes a truncated Gpr84 protein with two transmembrane domains deleted, likely rendering a non-functional GPCR. In WT zebrafish larval spinal cord at 6 days post-fertilization (dpf), astrocyte cell bodies are adjacent to the midline and emanate fine processes to infiltrate the neuropil, forming a dense meshwork in the lateral regions (Chen *et al*., 2020). Using the astrocyte transgenic line *Tg(slc1a3b:myrGFP-P2A-H2AmCherry)*, in which astrocyte membranes are labeled with myrGFP and nuclei labeled with H2AmCherry, we examined *gpr84^vo87/vo87^* homozygous mutants in comparison with the control siblings. Astrocytes in *gpr84^vo87/vo87^* homozygous mutants looked grossly normal at 6 dpf (Figure S5C), suggesting *gpr84* is not required for astrocyte development in zebrafish.

We next analyzed single-cell RNA-seq and Brain RNA-seq data in zebrafish and mammals (Farnsworth et al., 2020; Spanjaard et al., 2018; Zhang et al., 2014; Zhang et al., 2016) to identify genes encoding GPCRs, especially lipid-binding GPCRs, that are highly enriched in astrocytes. The gene *s1pr1* (*sphingosine-1-phosphate receptor 1*), which encodes a phospholipid-binding GPCR and belongs to the same GPCR family as *Drosophila* Tre1 (Figure S5A), showed noticeably higher expression in astrocytes. Interestingly, the S1pr1 GPCR has been shown to regulate germ cell migration in the ascidian *B. schlosseri* (Kassmer *et al*., 2015), similar to the role of Tre1 GPCR in *Drosophila* germ cells (Kunwar *et al*., 2003). In addition, *in vitro* studies have suggested that S1pr1 can regulate astrocyte morphology in mammals (Singh et al., 2021). By whole-mount *in situ* hybridization, we validated that *s1pr1* was expressed in zebrafish larval brain and spinal cord at 3 dpf (Figure S5D). We then used CRISPR/Cas9 to generate two independent *s1pr1* mutant alleles, *s1pr1^vo88^* and *s1pr1^vo89^*, which contain a 270 base-pair or 277 base-pair deletions, respectively, in the coding region of zebrafish *s1pr1* (Figure S5E). Interestingly, we found that astrocyte infiltration to the spinal cord neuropil in the lateral regions was severely disrupted in *s1pr1^vo88/vo88^* homozygous mutants in comparison with controls at 6 dpf (Figure 5A). In the *Tg(slc1a3b:myrGFP-P2A-H2AmCherry)* transgenic background, myrGFP-labeled dense astrocyte membranes in the lateral regions of the spinal cord were dramatically diminished in *s1pr1^vo88/vo88^* mutants, while the H2AmCherry-labeled nuclei number and position remained largely unaffected. These data argue that loss of S1pr1 alters astrocyte morphology, but not cell number. Comparable astrocyte defects were observed in *s1pr1^vo88/vo89^* trans-heterozygous mutants (Figure S5F), and we further validated the phenotypes with the astrocyte marker anti-GS to label zebrafish astrocytes and observed similar results (Figure S5G).

**Figure 5.**
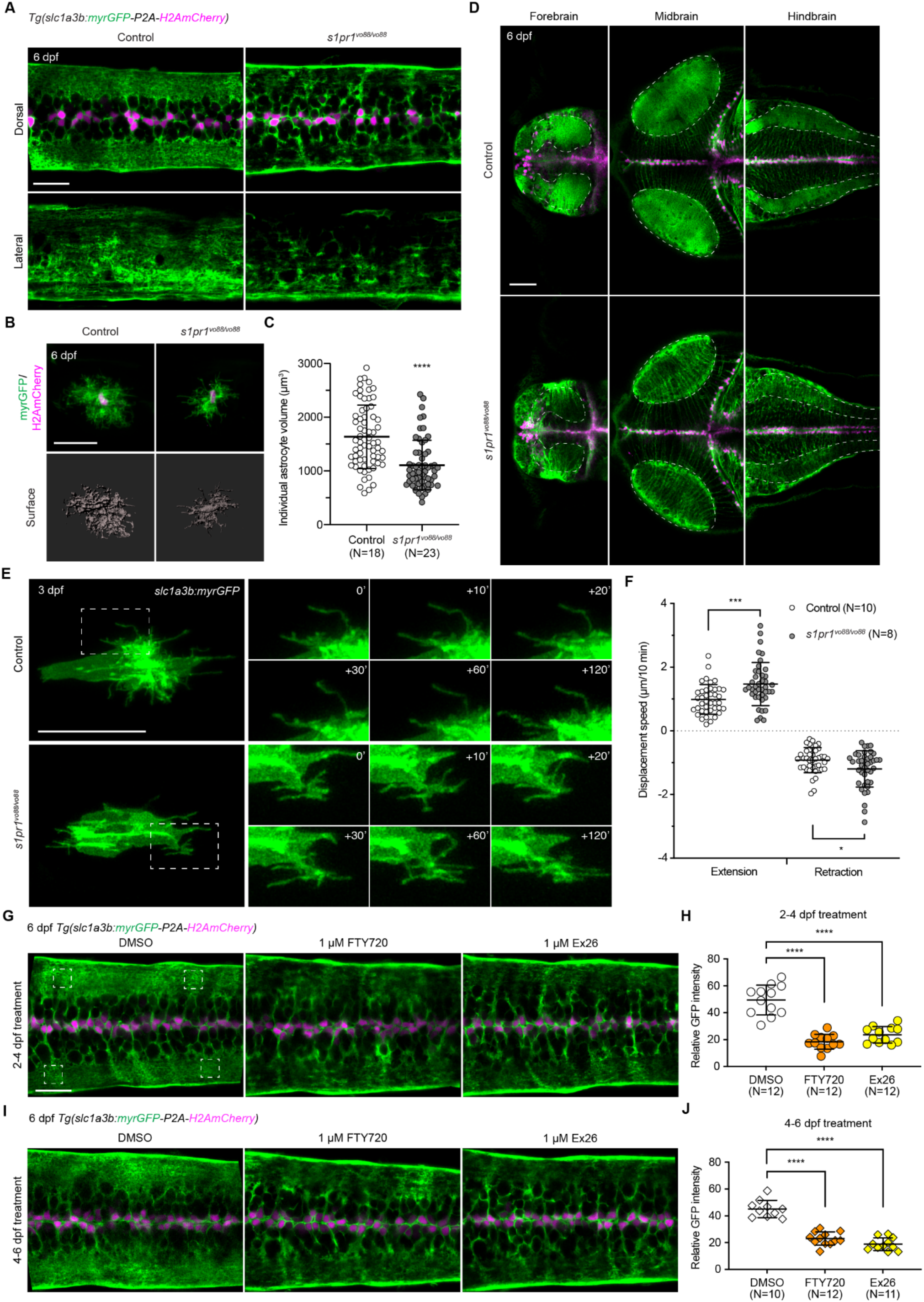
S1pr1 regulates astrocyte morphogenesis in zebrafish. (A) Images of spinal cord astrocyte membrane labeled with myrGFP (green) and nuclei labeled with H2AmCherry (magenta) in 6 dpf *Tg(slc1a3b:myrGFP-P2A-H2AmCherry)* transgenic control (N=15, dorsal view; N=3 lateral view) and *s1pr1^vo88/vo88^* mutants (N=17, dorsal view; N=5, lateral view). N, number of animals. Scale bar, 20 µm. (B) Images of sparsely labeled individual astrocytes by *slc1a3b:myrGFP-P2A-H2Amcherry* DNA constructs and the representative IMARIS 3D-rendering surface (grey) at 6 dpf in the spinal cord of control and *s1pr1^vo88/vo88^* mutant zebrafish. Scale bar, 20 µm. (C) Quantification of individual astrocyte volumes in control (N=18) and *s1pr1^vo88/vo88^* mutants (N=23) at 6 dpf. N, number of animals. Data points represent single astrocytes. ****, p<0.0001; unpaired t test. Error bars, mean values ± S.D. (D) Images of 6 dpf *Tg(slc1a3b:myrGFP-P2A-H2AmCherry)* larval brain in control (N=5) and *s1pr1^vo88/vo88^* mutants (N=6). Dashed lines mark astrocyte processes densely infiltrated regions, which are likely synapse-enriched neuropil as well, in the forebrain, midbrain, and hindbrain. Scale bar, 50 µm. (E) Time-lapse still images of astrocyte process dynamics labeled with myrGFP (green) in control and *s1pr1^vo88/vo88^* mutants at 3 dpf. Dashed boxes mark the regions shown to the right. Scale bar, 20 µm. (F) Quantification of astrocyte individual process extension and retraction displacement speed in control (N=10) and *s1pr1^vo88/vo88^* mutants (N=8) at 3 dpf. N, number of animals. Data points represent single astrocyte processes tracked. *, p<0.05; ***, p<0.001; unpaired t test. (G and I) Images of 6 dpf *Tg(slc1a3b:myrGFP-P2A-H2AmCherry)* transgenic larval spinal cord astrocytes after treatment with DMSO, 1 µM FTY720, or 1 µM Ex26 at 2-4 dpf (G) or 4-6 dpf (I). Dashed boxes represent 4 independent 10 µm x 10 µm areas in the astrocyte process-enriched regions were used to quantify the mean GFP intensity. Scale bar, 20 µm. (H) Quantification of relative GFP intensity at 6 dpf in the astrocyte process-enriched regions in DMSO (N=12), FTY720 (N=12), and Ex26 (N=12) after 2-4 dpf treatment. N, number of animals. Data points represent average GFP intensity of the 4 independent areas in a single fish. ****, p<0.0001; one-way ANOVA with multiple comparisons. (J) Quantification of relative GFP intensity at 6 dpf in the astrocyte process-enriched regions in DMSO (N=10), FTY720 (N=12), and Ex26 (N=11) after 4-6 dpf treatment. N, number of animals. Data points represent average GFP intensity of the 4 independent areas in a single fish.****, p<0.0001; one-way ANOVA with multiple comparisons. See also Figure S5 and Video S2.

To analyze astrocyte morphology at the single-cell level, we injected *slc1a3b:myrGFP-P2A-H2AmCherry* DNA constructs into one-celled zygotes to allow for sparse labeling and examined individual astrocyte clones at 6 dpf in the spinal cord. In contrast to control WT astrocytes, which exhibited a ramified morphology with many fine processes, individual *s1pr1^vo88/vo88^* mutant astrocytes occupied a smaller spatial domain and had significantly reduced volume sizes (Figures 5B and 5C). Astrocyte development was similarly perturbed throughout the CNS, including in in the forebrain, midbrain, and hindbrain of 6 dpf zebrafish larvae (Figure 5D), suggesting a requirement for S1pr1 in astrocyte morphogenesis throughout the CNS. Given that Tre1 and S1pr1 are both lipid-binding GPCRs and have been implicated in germ cell migration (Kassmer *et al*., 2015; Kunwar *et al*., 2003), our results argue that S1pr1 is the functional GPCR analog of Tre1 in vertebrate astrocytes.

### S1pr1 functions to balance astrocyte process outgrowth dynamics and is required throughout development

We showed previously that zebrafish astrocytes acquire their complex morphologies between 2 and 4 dpf in the larval spinal cord (Chen *et al*., 2020). We therefore used *slc1a3b:myrGFP* to sparsely label single astrocytes and performed time-lapse confocal microscopy to monitor fine cellular process dynamics at 3 dpf. We observed that astrocyte processes in control astrocytes displayed consistent extension and retraction rates throughout their morphogenesis (Figures 5E and 5F; Video S2). However, *s1pr1^vo88/vo88^* mutant astrocyte processes showed both faster extension and retraction speeds in comparison with controls, leading to a net reduction in stabilized astrocyte processes compared to controls (Figures 5E and 5F; Video S2). These data demonstrate that astrocyte membrane process dynamics are disrupted in *s1pr1* mutants during astrocyte growth.

S1pr1 modulators, such as FTY720 and Ex26, can be used to acutely block S1pr1 signaling *in vivo* (Cahalan et al., 2013; Matloubian et al., 2004). We used these reagents to determine the temporal requirements for S1pr1 signaling in astrocyte morphogenesis. We first treated *Tg(slc1a3b:myrGFP-P2A-H2AmCherry)* transgenic WT zebrafish with 1 µM FTY720 or 1 µM Ex26 from 2 to 4 dpf, stages when astrocytes are rapidly growing in the spinal cord and found that inhibiting S1pr1 by both FTY720 and Ex26 between 2-4 dpf resulted in disrupted astrocyte morphology by 6 dpf (Figure 5G). Quantification of GFP mean intensity in astrocyte process-enriched regions revealed that the astrocyte membrane signal was significantly reduced in FTY720/Ex26-treated larvae compared to DMSO-treated controls (Figure 5H). To determine whether such effects were mediated specifically through S1pr1, we performed similar treatment experiments in the *s1pr1^vo88/vo88^* mutant backgrounds between 2-4 dpf, and we found that the astrocyte infiltration defects in *s1pr1^vo88/vo88^* mutants were not further exacerbated by either FTY720 or Ex26 (Figures S5H and S5I).

We next asked if S1pr1 inhibition could alter the morphology of more mature astrocytes. We treated zebrafish larvae with FTY720 or Ex26 between 4-6 dpf before imaging. Interestingly, treatment of *Tg(slc1a3b:myrGFP-P2A-H2AmCherry)* transgenic WT larvae also resulted in disruption of astrocyte infiltration of the synaptic neuropil (Figures 5I and 5J). Notably, GFP mean intensity quantification demonstrated that the astrocyte defects in 4-6 dpf treatment experiments were comparable to that observed in 2-4 dpf treatment experiments (Figures 5H and 5J). Finally, we did not observe exacerbated astrocyte phenotypes for 4-6 dpf FTY720/Ex26 treatment experiments in the *s1pr1^vo88/vo88^* mutants, implying the late requirement in astrocyte morphogenesis is also S1pr1-dependent (Figures S5J and S5K). Based on these results, we conclude that S1pr1 GPCR signaling is required throughout astrocyte growth for normal elaboration and maintenance of astrocyte morphology *in vivo*.

### Wunens play opposing roles in astrocytes and neurons to influence astrocyte infiltration

In *Drosophila*, lipid phosphate phosphatases (LPP) Wunen (Wun) and Wunen2 (Wun2) have been shown to function redundantly to hydrolyze phospholipids and to promote the uptake of lipid product into cells (Renault et al., 2004; Starz-Gaiano et al., 2001). Similarly, the vertebrate homologues of Wunen, LPP1-3, can regulate extracellular S1pr1 ligand sphingosine-1-phosphate (S1P) levels and its activity (Tang et al., 2015). Loss of *wun*/*wun2* results in germ cell migration defects, which argues that Wun/Wun2 also act upstream of Tre1 in *Drosophila* (Kunwar *et al*., 2003; LeBlanc and Lehmann, 2017). We therefore sought to test the role of Wunens/LPPs in astrocyte morphogenesis.

We first analyzed astrocyte morphology using different *wun* or *wun2* mutants over a deficiency that deletes both *wun/wun2*. Strong *wun*/*wun2* double mutants resulted in pupal lethality, so we analyzed mutant phenotypes in wandering third instar larvae (wL3). Using anti-Gat to label astrocyte membranes in the ventral nerve cord (VNC), we found that loss of single *wunen* genes (*wun^23^*, *wun^9^*, or *wun2^N14^*) did not alter astrocyte morphology in comparison with controls (Figure S6A). However, loss of both *wun* and *wun2* (*wun^49^,wun2^EX34^*) together resulted in astrocyte infiltration defects in the VNC (Figure S6A). This is consistent with previous observations that Wun and Wun2 can act redundantly in other tissues (Renault *et al*., 2004). To determine whether the astrocyte infiltration phenotypes in *wun*/*wun2* mutants were due to a cell-autonomous requirement for Wunen activity, we used the astrocyte driver *alrm-Gal4* to express *wun* or *wun2* specifically in astrocytes. We found that expression of *wun2* or *wun* was sufficient to rescue the infiltration defects to levels similar to controls, while a catalytically dead mutant construct (*wun2^H326K^*) failed to rescue astrocyte infiltration defects in *wun^49^,wun2^EX34^* mutant backgrounds (Figures 6A and 6B). Overexpression of *wunens* (*wun2* or *wun*) in astrocytes in a control background using the *alrm-Gal4* driver or a stronger astrocyte driver (*GMR25H07-Gal4*) did not alter astrocyte morphology (Figure S6B). In addition, astrocyte expression of mouse *LPP3* (*mLPP3*) in *wun^49^,wun2^EX34^* mutant animals was able to suppress the infiltration defects (Figures 6A and 6B), demonstrating that mLLP3 can substitute for Wunen activity in *Drosophila* astrocytes.

**Figure 6.**
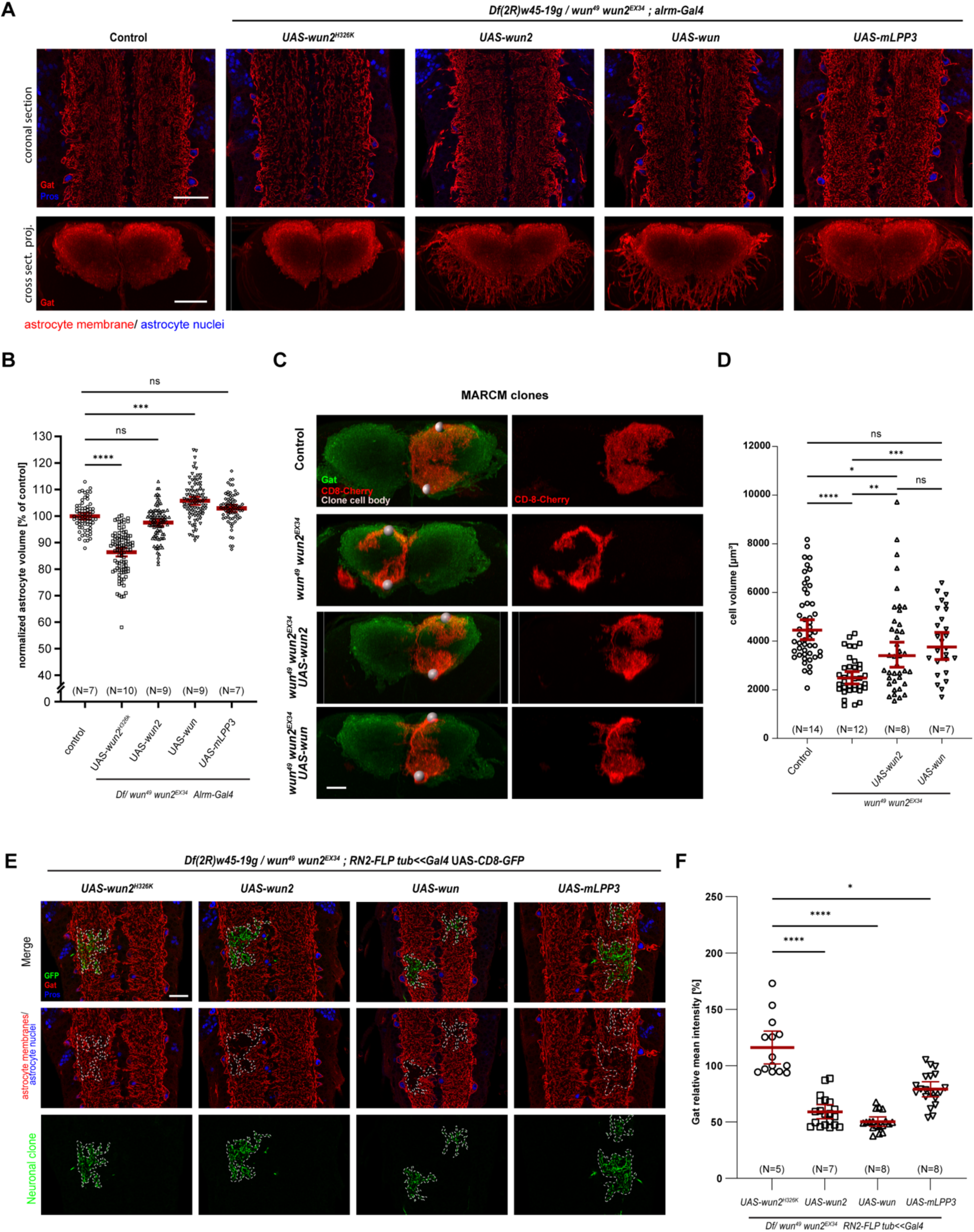
The *Drosophila* LPPs Wun/Wun2 control astrocyte infiltration. (A) Confocal single section images of ventral nerve cords of third instar larvae (L3) in coronal sections (upper panels) or cross-sectional 3D projections of 100 µm along the anterior-posterior axis. Astrocyte membranes are labeled by α-Gat antibody (red) and nuclei labeled by α-Pros antibody (blue). Scale bars, 30 µm. (B) Quantification of normalized astrocyte infiltration densities of control (*w^1118^*, N= 7); and *Df(2R)w45-19g* / *wun49 wun2^Ex^*^34^; *alrm*-Gal4/+ animals expressing UAS-*wun2^H326K^* (N= 10), UAS-*wun2* (N= 9), UAS-*wun* (N= 9), UAS-*mLPP3* (N= 7) related to (A). N denotes the number of animals, and individual data points represent astrocyte volume measured in a small sampling volume of 200*100*20 pixels, with five volumes per left and five volumes per right side of the mid-dorsal neuropil region along the a-p axis totaling 10 measurements per larva. Kruskal-Wallis test with Dunn’s multiple comparisons test. Error bars: mean values ± 95% CI. ns, not significant; *, p<0.05; **, p<0.01; ***, p<0.001; ****, p<0.0001. (C) Cross-sectional 3D projections of MARCM astrocyte clones labeled with mCD8-mCherry (red) and all astrocyte membrane labeled with α-Gat antibody (green) in the L3 VNC. Cell bodies of the MARCM astrocyte clones are indicated with grey spheres for clarity. Scale bar, 15 µm. (D) Quantification of astrocyte volumes of MARCM clones for control (N= 14); *wun^49^*,*wun2^EX34^* (N= 12); *wun^49^*,*wun2^EX34^* + UAS-*wun2* (N= 8); and *wun^49^*,*wun2^EX34^* + UAS-*wun* (N= 7) related to (C). N denotes the number of animals and individual data points represent single clones. Kruskal-Wallis test with Dunn’s multiple comparisons test. Error bars: mean values ± 95% CI. ns, not significant; *, p<0.05; **, p<0.01; ***, p<0.001; ****, p<0.0001. (E) Images of L3 VNC with astrocyte membranes labeled with α-Gat (red) and astrocyte nuclei labeled with α-Pros (blue) in the *Df(2R)w45-19g* /*wun^49^*,*wun2^EX34^; RN2-FLP tub<<Gal4* UAS-CD8-GFP FLP-out background expressing different *wunen* or *mLPP3* transgene in the mCD8-GFP-labeled neuronal clones (green). Dashed lines mark the boundaries of mCD8-GFP-labeled neurons at a single-Z plane. Scale bar: 20 µm. (F) Quantification of relative α-Gat mean fluorescence intensity in the dashed line areas normalized to neighboring regions of the same size lacking a neuronal clone related to (E). UAS-*wun2^H326K^* (N= 5); UAS-*wun2* (N= 7); UAS-*wun* (N=8); UAS-*mLPP3* (N= 8); Kruskal-Wallis test with Dunn’s multiple comparisons test. Error bars: mean values ± 95% CI. *, p<0.05; ****, p<0.0001. See also Figure S6.

Unexpectedly, we observed that astrocyte processes extended out of the synaptic neuropil and into the cortex when Wunen activity was restored in astrocytes (Figure 6A). Astrocyte membranes normally remain restricted to the synaptic neuropil unless cortex glia are ablated (Coutinho-Budd et al., 2017). To further explore how Wunen regulates astrocyte growth, we employed the Mosaic Analysis with a Repressible Cell Marker (MARCM) approach (Lee and Luo, 1999) to analyze *wun^49^,wun2^EX34^* astrocyte mutant clones. Compared to control clones, we found that similar to *Tre1* mutants, individual mutant astrocyte volumes of *wun/wun2* mutant clones were significantly reduced (Figures 6C and 6D). When *wun2* or *wun* was re-expressed in the mutant MARCM clones, the overall astrocyte morphology and cell volumes were significantly rescued (Figures 6C and 6D). Together, these data indicate that Wunen activity is required in astrocytes for normal morphology.

The ectopic astrocyte processes we observed when overexpressing single Wunen in the *wun/wun2* mutants but not in a WT background (Figures 6A and S6B) suggest Wunen might play a role in regulating cell-cell interactions and/or competition with surrounding cells. To explore this further, we first used a pan-neuronal driver *n-syb-Gal4* (Rao et al., 2001) to misexpress *wun* in all neurons and examined astrocyte infiltration phenotypes. In a WT background, we found that Wunen activity in neurons only moderately altered astrocyte infiltration in the VNC (Figure S6C). In contrast, neuronal expression of *wun* in the *wun^49^,wun2^EX34^* mutant background led to dramatic reduction in astrocyte infiltration of the neuropil, which was not replicated when we used the catalytically dead mutant *wun2^H326K^* (Figure S6C). We next used a FLP-out method with *RN2-FLP tub<<Gal4* (Ou et al., 2008) to manipulate Wunen levels in a subset of mCD8-GFP-labeled neuronal clones in the VNC. Clonal misexpression of Wun in single neurons in a WT background led to a mild reduction of astrocyte processes within the area of the neuronal clone (Figure S6C). However, we found that expression of *wun* or *wun2* in single neurons resulted in a near complete elimination of astrocytic processes from the domain occupied by the *wunen*-expressing neuron in the *wun/wun2* mutant background (Figures 6E and 6F). This phenotype depended on Wun enzymatic activity and could also be observed, to a reduced extent, after *mLPP3* expression (Figures 6E and 6F). We conclude that neuronal misexpression of Wunen can block the morphological elaboration of surrounding astrocytes, and our data suggests that Wunen activity likely functions very locally in regulating astrocyte growth.

In conclusion, we have shown that Wunen activity is required cell autonomously in astrocytes to promote their proper process elaboration. In addition, we find that Wunen expression in neurons can block astrocyte process outgrowth, most likely by depleting an attractive, growth promoting phospholipid in the environment.

### Wunen regulation of astrocyte growth depends on Tre1

We used anti-Gat to label astrocyte membranes to compare the astrocyte morphology in control animals, *wun^49^,wun2^EX34^* mutants, *Tre1^attP^* mutants, and the *Tre1^attP^*;*wun^49^,wun2^EX34^* triple mutants. As described above, we observed mild astrocyte infiltration defects in *wun^49^,wun2^EX34^* mutants, but stronger infiltration defects in *Tre1^attP^* mutants (Figure 7A). When we compared *Tre1^attP^*; *wun^49^,wun2^EX34^* triple mutants with *Tre1^attP^* mutants, we found they showed similar astrocyte infiltration defects (Figure 7A). This non-additivity suggests that Tre1 and Wun/Wun2 are in the same genetic pathway in regulation of astrocyte growth. To determine whether Wunen-dependent regulation of astrocyte growth requires Tre1, we used a FLP-out approach to sparsely express *wun2* in myr::tdTomato-labeled astrocyte clones in *wun^49^,wun2^EX34^* mutants or *Tre1^attP^*;*wun^49^,wun2^EX34^* triple mutants. In the *wun^49^,wun2^EX34^* mutant background, *wun2*-expressing astrocyte clones displayed dramatically increased volumes and expanded their territories broadly across the neuropil (Figures 7B-D). This observation further supports the notion that Wunens can potently regulate astrocyte growth. In contrast, in the *Tre1^attP^*; *wun^49^,wun2^EX34^* triple mutant background, we found that Wunen-mediated astrocyte overgrowth phenotypes were completely suppressed (Figures 7B-D). The expanded process outgrowth we observed in Wun2-overexpressing astrocyte clones in the *wun^49^,wun2^EX34^* mutant background was restricted to the neuropil without prominent projections entering the cortex (Figure 7B), contrary to what might have been expected from global astrocyte expression (Figure 6A). However, clone-adjacent astrocytes (but not those distant from clones) frequently showed prominent membrane projections into the cortex (Figure 7B, arrowheads). The formation of these ectopic astrocyte processes was completely suppressed in a *Tre1^attP^*;*wun^49^,wun2^EX34^* triple mutant background, indicating this non-cell autonomous effect of Wun2-overexpressing clones is also dependent on Tre1 activity. Together, these data argue strongly that Wunens act in the same signaling pathway as Tre1 to regulate astrocyte morphogenesis, that Tre1 acts downstream of Wunens, and that astrocytes compete with their neighbors for an attractive growth promoting phospholipid that can be inactivated by LPPs.

**Figure 7.**
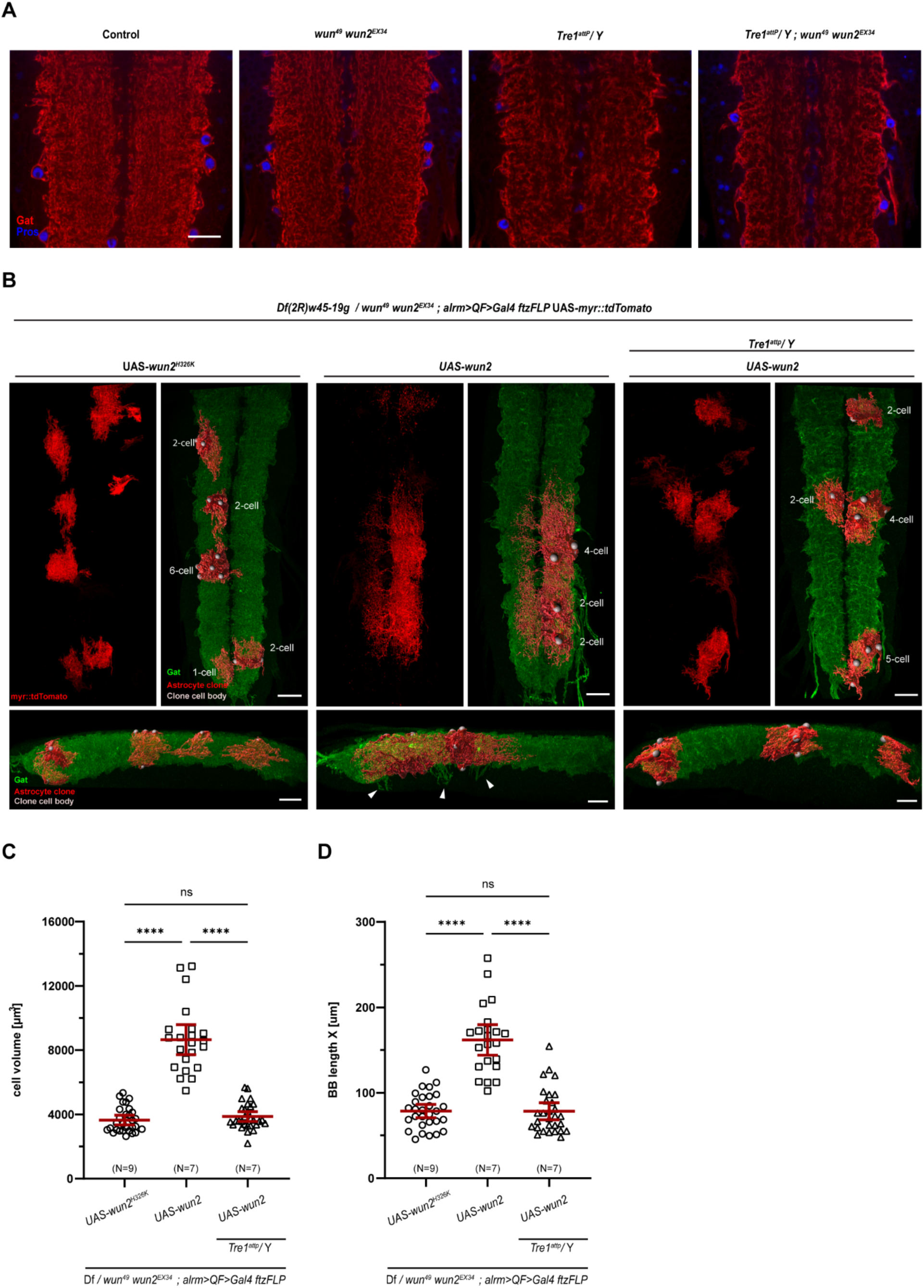
Wunen activity is suppressed by the loss of *Tre1*. (A) Confocal single section images of L3 VNCs with astrocytes membranes labeled by α-Gat antibody (red) and nuclei labeled by α-Pros antibody (blue) in control (N= 9) or *wun^49^*,*wun2^EX34^* (N= 7), *Tre1^attP^* (N= 14), and *Tre1^attP^*;*wun^49^*,*wun2^EX34^* (N= 12) mutants. N denotes the number of animals and individual data points represents single clones. Scale bar, 20 µm. (B) 3D projections of confocal images of astrocyte FLP-out clones labeled with myr::tdTomato (red) using *alrm*>QF>Gal4 *ftz*-FLP with all astrocytes labeled by α-Gat (green) in the L3 VNC. Clones also express UAS-*wun2^H326K^* or UAS-*wun2* in either the *Df(2R)w45-19g/*/*wun^49^*,*wun2^EX34^* mutant background or the *Tre1^attP^*; *Df(2R)w45-19g/*/*wun^49^,wun2^EX34^* triple mutant background. The right and bottom panels shows 3D renderings of segmented astrocyte clones, and cell bodies of cells in the clone are highlighted with grey spheres for clarity. Bottom panels show a lateral view of the VNC. Arrowheads point out ectopic projections into the cortex of astrocytes adjacent to the astrocyte clones expressing UAS-*wun2*. Scale bars, 30 µm. (C and D) Quantification of the cell volume of individual astrocytes (C) and the length of a minimal bounding box (BB) enclosing astrocyte clones (D) in UAS-*wun2*-expressing astrocytes (related to (B): *Df(2R)w45-19g*/*wun^49^*,*wun2^EX34^* + UAS-*wun2^H326K^* (N= 9); *Df(2R-w45-19g*/*wun^49^*,*wun2^EX34^* + UAS-*wun2* (N= 7); and *Tre1^attP^/Y*; *Df(2R)w45-19g*/*wun^49^,wun2^EX34^* + UAS-*wun2* (N= 7). N denotes the number of animals and individual data points represents single clones. Kruskal-Wallis test with Dunn’s multiple comparisons test. Error bars: mean values ± 95% CI. ns, not significant; ****, p<0.0001.

## DISCUSSION

Our work demonstrates the Tre1/S1pr1 GPCR signaling pathway and the lipid phosphate phosphatases (LPPs) Wun and Wun2 are essential for astrocyte morphogenesis in *Drosophila* and zebrafish. Although *s1pr1* is not the closest sequence homolog of *Tre1* in vertebrates, both S1pr1 and Tre1 belong to the rhodopsin-like GPCR family and share the same highly conserved signaling motifs (Zhou et al., 2019). In addition, these phospholipid-binding GPCRs and S1P have been shown to functionally interact with LPPs in analogous ways during germ cell migration (Kassmer *et al*., 2015; LeBlanc and Lehmann, 2017; Paksa et al., 2016). Given the strikingly similar astrocyte phenotypes we observed between *Drosophila Tre1* mutants and zebrafish *s1pr1* mutants (Figures 1 and 5) and the fact that they share the same ligand processing enzymes, we propose that Tre1/S1pr1 represent a functionally conserved phospholipid-mediated GPCR signaling pathway that regulates astrocyte development.

There are important phenotypic differences between the roles of Tre1 and Wun/Wun2 in germ cells versus astrocytes. In germ cells, Tre1 and Wun/Wun2 guide germ cell migration to the gonad and also regulate germ cell survival (Hanyu-Nakamura et al., 2004; Kunwar *et al*., 2003; LeBlanc and Lehmann, 2017; Renault *et al*., 2004). In astrocytes, we show these molecules regulate astrocyte process outgrowth and establishment of morphological complexity. We have found no evidence for a role for Tre1/S1pr1 or Wun/Wun2 in astrocyte survival. Our data support the notion that Tre1 functions to balance Rac1 activity through its NPIIY motif to help organize the astrocyte cytoskeleton (Figures 3 and 4). Rac1 has a well-established role in actin cytoskeleton regulation in development and disease (Liang et al., 2021; Luo *et al*., 1994), and Rac1 is directly involved at the cell membrane to remodel actin dynamics for membrane protrusive behaviors (Ridley et al., 1992). Live-imaging in zebrafish revealed a role for S1pr1 in the earliest stages of astrocyte process outgrowth, with *s1pr1* mutant astrocytes exhibiting much more dynamic membrane extension/retraction (Figures 5E and 5F). The simplest interpretation of our data is that Tre1/S1pr1 GPCR signaling orchestrates Rac1 activity at the plasma membrane to influence actin-based cellular process protrusions in astrocytes. How could changes in protrusion dynamics lead to a simplified astrocyte morphology? In mouse and zebrafish studies, S1pr1 acts as a negative regulator in endothelial cell growth, by preventing excessive sprouting during angiogenesis and vessel maturation, and the mature vascular network in *s1pr1* mutants is ultimately disorganized with enlarged vessel diameter and reduced vessel number (Ben Shoham et al., 2012). It is possible that excessive extension/retraction of astrocyte processes in *Tre1* or *s1pr1* mutants similarly disrupts astrocyte morphology, perhaps by altering the ability of astrocytes to form adhesive interactions with surrounding cells, ultimately leading to a simplification of fine processes.

The precise nature of the extracellular phospholipids that activate Tre1 or S1pr1 signaling *in vivo* has been difficult to resolve. To our knowledge, ligands for Tre1 have not been identified. S1P, which is speculated to activate S1pr1 and regulate germ cell migration (Kassmer *et al*., 2015; Mendelson et al., 2014), is a candidate for regulating astrocyte growth in vertebrates. The notion that germ cell migration is driven by lipids was first shown in *Drosophila*, where *wun* and *wun2* mutants were found to perturb germ cell migration to the gonad (Zhang *et al*., 1997), and other lipid-modifying enzymes have since been shown to play a role in germ cell guidance (Richardson and Lehmann, 2010). We have shown that Wun and Wun2 are potent regulators of astrocyte growth, and this depends on their LPP enzymatic activity. These observations provide strong support for the notion that extracellular phospholipids modulate astrocyte membrane process growth. Where these bioactive phospholipids come from, and how many can regulate astrocyte growth, are open questions, as is the precise nature of their signaling to astrocytes.

How do Tre1 and Wun/Wun2 regulate growth? We found that Wun/Wun2 cell-autonomously facilitate Tre1-dependent growth of astrocytes while neuronal expression of these LPPs can suppress astrocyte outgrowth within the domain of Wun-expressing neurons. The simplest interpretation of our data, is that extracellular phospholipids activate Tre1 to positively drive astrocyte growth, and that Wun/Wun2 act locally to process this ligand (Figure S7). We can envision a number possible mechanisms by which Tre1 and Wun/Wun2 signaling might occur. First, after phospholipid binding to Tre1, the activated receptor promotes astrocyte growth, is internalized, and Wunens dephosphorylate the phospholipid ligand bound to Tre1 to enable receptor recycling to the plasma membrane and continued Tre1 signaling. Interestingly, downregulation of surface S1pr1 receptors is how the therapeutic inhibitor fingolimod is believed to block S1pr1 signaling (Matloubian *et al*., 2004). Second, Wunens could dephosphorylate phospholipids outside the cell, thereby allowing entry, where lipids are rephosphorylated by sphingosine kinase to activate Tre1 in intracellular vesicles. This is supported by data showing that Wun/Wun2 can promote lipid entry into cells (Kono et al., 2022; Renault *et al*., 2004). Third, after entry of dephosphorylated lipids into the cell, lipids might be rephosphorylated in a way that makes them bioactive for Tre1, then be transported out of the cell where they can locally activate Tre1 receptors (Kono *et al*., 2022). Ultimately, astrocyte growth would be driven by astrocyte-astrocyte competition for these growth-promoting extracellular phospholipids. According to this model, the ability of neuronal expression to block astrocyte growth would be due to the ability of Wunens to degrade this growth promoting phospholipid, in the domain of the neuron expressing LPPs. Likewise, the lack of overgrowth when Wun or Wun2 are expressed in control animals (versus a *wun, wun2* mutant) would be explained by the ability of neighboring cells to complete for phospholipids.

*Tre1* mutant astrocyte phenotypes are more pronounced than those seen in *wun,wun2* double mutants. These observations may indicate that additional LPPs regulate extracellular phospholipid levels involved in astrocyte growth, or other LPP-independent ligands signal through Tre1 in *Drosophila*. Tre1 signaling appears to be sensitive to the relative levels of phospholipids, and Wun/Wun2 seem to regulate the spatial profile of these extracellular lipids. For instance, while clonal Wun or Wun2 overexpression in a WT background did not dramatically alter astrocyte morphology, expression of Wun2 in astrocytes in a *wun,wun2* double mutant background resulted in massive overgrowth of astrocytes in a Tre1-dependent fashion. Here, the Wun2-expressing cell has a competitive advantage over its neighbors due to its ability to facilitate Tre1 signaling cell autonomously and the inability of the neighboring cells to inactivate the phospholipid in their territory.

How astrocytes determine their territories in relation to other astrocytes is poorly understood (DeSantis and Smith, 2021; Stork *et al*., 2014; Torres-Ceja and Olsen, 2022). Our data supports a model in which Tre1/S1pr1 and Wunen/LPP are well poised to control competitive interactions between neighboring astrocytes, analogous to the role in germ cell development (Renault et al. 2004). Tre1/S1pr1 acts as the receptor for a bioactive phospholipid to promote astrocyte process outgrowth and elaboration, and Wunen/LPP cell-autonomously facilitate Tre1/S1pr1 signaling while also degrading free phospholipids to deprive neighboring cells from this growth promoting factor. We propose that the Tre1/S1pr1 and Wunen/LPP system, together with other signaling systems like FGF-signaling (Chen *et al*., 2020; Kang et al., 2014; Stork *et al*., 2014) and cell contact dependent interactions (Baldwin et al., 2021a; Stogsdill *et al*., 2017; Takano et al., 2020), is a major contributor to astrocyte-astrocyte territorial competition and tiling (Bushong et al., 2002, 2004). Unexpectedly, in Wunen/LPP rescue experiments we observed prominent projections of astrocytes into the cortex (Figure 6A), suggesting that Wunen function is not restricted to astrocyte-astrocyte interactions but that astrocytes are also competing with cortex glia that ensheath the neuronal cell bodies (Coutinho-Budd *et al*., 2017). This not a strictly cell-autonomous function of Wun2 overexpression: clonal rescues show that while the Wun2 expressing cells show substantial overgrowth in the neuropil, it is the non-expressing immediately neighboring astrocytes that grow ectopic processes into the cortex (Figure 7B). This is likely due to the efficient depletion of active phospholipid in the neuropil by the Wun2-expressing clone, with residual phospholipid in the cortex that can still activate Tre1-dependent process outgrowth.

Our experiments to pharmacologically inhibit S1pr1 indicate that S1pr1 activity is continuously required to guide astrocyte morphogenesis during development at the larval stages in zebrafish (Figures 5G-J and S5H-K). It remains to be determined how astrocytes in the adult CNS respond to S1pr1 blockade, but this is an important question in the context of human health as the S1pr1 modulator FTY720 (Fingolimod) is a bioactive compound that has been used in humans to treat multiple sclerosis (MS) (Brinkmann et al., 2010; Kappos et al., 2006). FTY720 is thought to act through peripheral immune-based mechanisms to reduce egress of lymphocytes from lymph nodes to prevent T cell-elicited neuroinflammation to the CNS. FTY720 can also cross the blood-brain barrier to target astrocytes to ameliorate astrogliosis-associated neurodegeneration in MS (Choi et al., 2011). However, owing to the profound astrocyte morphological changes we observed in FTY720-treated zebrafish larvae, our findings raise the concern that FTY720 treatment might dramatically alter astrocytic morphology.

## ACKNOWLEDGMENTS

We would like to thank Dr. Y. Rao, Dr. R. Lehmann, Dr. A. Renault, and Dr. T. Awasaki for kindly providing us plasmids and flies, the Bloomington Stock Center and the Vienna Drosophila RNAi Center for fly stocks, and the Developmental Studies Hybridoma Bank, created by the NICHD of the NIH and maintained at The University of Iowa, Department of Biology, Iowa City, IA 52242 for select monoclonal antibodies. We thank A. Forbes and T. Perry for excellent zebrafish care, and colleagues from the Freeman and Monk labs for helpful comments and discussion on the manuscript. This work was supported by R37NS053538 to M.R.F., R21NS115437 to K.R.M., and R01NS124146 to M.R.F. and K.R.M.

## AUTHOR CONTRIBUTIONS

J.C., T.S., K.R.M., and M.R.F. conceived and designed the research. J.C., T.S., Y.K., and A.S. performed the experiments in *Drosophila*, and J.C. and C.P. performed the experiments in zebrafish. J.C., T.S., K.R.M., and M.R.F. wrote the manuscript, and all authors contributed to the final draft.

## METHODS

### Experimental model and subject details

#### Drosophila strains

Flies (*Drosophila melanogaster*) were kept on standard cornmeal agar with a 12hr/12hr light cycle at 25°C. The following lines were used: *w^1118^*, *Tre1^attP^* (Deng *et al*., 2019), *Tre1*^D^*^EP5^* and *Tre1^sctt^* (gift from R. Lehmann, Whitehead Institute, Cambridge, MA United States; LeBlanc and Lehmann, 2017), *Df(2R) BSC408* (Cook et al., 2012), *wun2^N14^*, *Df(2R)w45-19g* (Hanyu-Nakamura *et al*., 2004), *wun^23^*, *wun^9^* (Renault *et al*., 2004), *wun^49^ wun2^EX34^* (Renault et al., 2010), *UAS-wun (Zhang et al., 1997)*, *UAS-wun2-GFP* (Ile et al., 2012), and *UAS-wun2^H326K^* (Starz-Gaiano *et al*., 2001), *UAS-mLPP3* (Ile *et al*., 2012), *alrm-Gal4* (Doherty et al., 2009), *UAS-mCD8-GFP* (Lee and Luo, 1999), *tre1^Gal4^* (this study), *hsFLPD5.fco* (Nern et al., 2011), *alrm>nlsLexAfl>Gal4co* (this study), *alrm>QF>Gal4* (this study), *10XUAS-IVS-myrGFP* (Pfeiffer et al., 2010), 10XUAS-IVS-myr::tdTomato (Bloomington 32221), *UAS-Lifeact-GFP.W* (Bloomington, 57326), *UAS-chRFP-Tub* (Bloomington, 25773), *UAS-mCD8-mCherry* (Stork *et al*., 2014), *UAS-lam-GFP* (Aza-Blanc et al., 2000), *GMR25H07-Gal4* (Bloomington, 49145), *GMR25H07-LexA* (Bloomington, 52711), *13xLexAop2-mCD8-GFP* (Pfeiffer *et al*., 2010), *UAS-Tre1^KK102307^* RNAi (VDRC, v108952), *UAS-Tre1^HMS00599^* RNAi (Bloomington, 33718), *5xUAS-Tre1*, *5xUAS-Tre1^NAY^*, *5xUAS-Tre1^AAIIY^*, and *5xUAS-Tre1^NAY,AAIIY^* (this study), *UAS-Rac1.N17* and *UAS-Rac1.W* (Luo *et al*., 1994), *RN2-FLP tub-Gal4 UAS-mCD8-GFP* (Ou *et al*., 2008), *n-syb-Gal4* (Rao *et al*., 2001).

### Zebrafish husbandry and transgenic lines

All zebrafish studies were performed in compliance with institutional ethical regulations for animal testing and research at Oregon Health & Science University (OHSU). Experiments were approved by the Institutional Animal Care and Use Committee of OHSU. Zebrafish were maintained at 28°C and fed with a combination of rotifer suspension, brine shrimp, and dry food (Gemma 75, 150, 300). Zebrafish embryos and larvae were raised at 28.5°C in petri dishes with embryo medium, and phenylthiourea (PTU, 0.004% final concentration) was used to reduce pigmentation after 24 hpf for live imaging experiments. The following lines were used in this study: *gpr84^vo87^*, *s1pr1^vo88^*, and *s1pr1^vo89^* (this study), and *Tg(slc1a3b:myrGFP-P2A-H2AmCherry)* (Chen *et al*., 2020).

### Method details

#### Generation of *Tre1* endogenous Gal4 reporter flies

To generate the *Tre1* endogenous Gal4 reporter line (*Tre1^Gal4^*), we first modified the *pBsk-attB-13687-Gal4* (gift from Y. Rao, Peking University, Beijing China; Deng *et al*., 2019) to replace the *13687* sequence with NheI-AgeI-NotI restriction enzyme sites. We subsequently PCR amplified the genomic DNA sequence spanning the coding region of *Tre1* to be inserted using NheI and AgeI to generate *pBsk-attB-Tre1-Gal4*. The construct was sequence verified by Sanger sequencing (Genewiz) and used for transgenesis (Bestgene), and the resulting stable stock was further crossed with a Cre line (Bloomington, 1092) to excise the exogenous markers (*w^+^* and *3xP3-RFP*) via the flanking *LoxP* sites (Figure S1). Primer sequences are described in Table S1.

#### Generation of *UAS-Tre1* expressing transgenes

We PCR amplified the sequences from constructs of *nosp-tre1^+^ flag*, *nosp-tre1 NRY^−^ flag*, *nosp-tre1 NPIIY^−^ flag*, and *nosp-tre1 NRY^−^NPIIY^−^ flag* (gift from R. Lehmann, Whitehead Institute, Cambridge, MA United States; LeBlanc and Lehmann, 2017) and subcloned into the *pattB-5xUAS* vector using XhoI and XbaI digestion sites.

All the constructs were sequence verified by Sanger sequencing (Genewiz) and used for *attP2* landing site-specific integration on the 3^rd^ chromosome. Primer sequences are described in Table S1.

#### Generation of *alrm*>QF>Gal4 (*alrm*-QNUG)

The *alrm*-QNUG (**Q**F **N**o **U**TR **G**al4co) was generated similar to a previously published version of this flip-out construct (Stork et al. 2014) but, lacks the dedicated 3’ Hsp70 terminator for the QF cassette but retains a SV40 terminator for the Gal4co. In short, QF was amplified from *pCaSpeR-EFAN-FRT-QF-HSP70-FRT-Gal4co* using the primers *QF NotI for* and *SpeI FRT QF noUTR rev* and cloned into *pCaSpeR-EFAN-Gal4co* via NotI/SpeI sites to form *pCaSpeR-EFAN FRT-QF-noUTR-FRT-Gal4co*. The *alrm* promotor was amplified from genomic DNA with the primers *Alrm FseI for* and *Alrm AscI rev* and inserted into *pCaSpeR-EFAN FRT-QF-noUTR-FRT-Gal4co* via FseI/AscI sites to form *pCaSpeR-EFAN-alrm-QNUG*.

#### Generation of *gcm*-FLP and *ftz*-FLP

A DNA fragment containing a DSCP basal promoter, Syn21 5’UTR (Pfeiffer et al., 2012) FLPD5 (Nern et al., 2011) coding sequence and a P10 3’UTR (Pfeiffer et al., 2012) were codon optimized and synthesized by Genescript and delivered in a *puc57* vector. Subsequently this fragment was transferred into a modified *pCaSpeR* vector *pCaSpeR-EFAN* (Stork et al. 2014) via NotI/XhoI sites to form *pCaSpeR-EFAN-FLP*.

The *gcm* promoter region of 9kb (Ragone et al., 2002) was amplified from the genomic P[acman] BAC clone CH322-71L23 (Venken et al., 2009) in two 4.5 kb fragments with the primers *gcm 9-4.5 Gibson for*, *gcm 9-4.5 rev*, *gcm 4.5-0 for* and *gcm 4.5-0 Gibson rev* that were subsequently cloned into an AscI/ NotI digested *pCaSpeR-EFAN-FLP* using Gibson assembly (Gibson et al. 2009). For the *ftz*-FLP construct, a 127 bp fragment of the *fushi tarazu* Promoter/5’UTR (Jacobs et al. 1989) was PCR amplified from genomic DNA with the primers *ftz for NotI* and *ftz rev BamHI* and cloned into *pCaSpeR-EFAN-FLP* via NotI/BamHI restriction removing the DSCP basal promoter from the final construct. The *alrm*>nlsLexAfl>Gal4co, *alrm*>QF>Gal4, *gcm*-FLP, and *ftz*-FLP transgenes were generated by P-element mediated transgenesis at Bestgene.

#### Generation of mutants in zebrafish

We used CRISPR/Cas9 method to generate genetic mutants in zebrafish. The web tool of CHOPCHOP (Labun et al., 2019) was used to select target sites, and individual *sgRNAs* were synthesized using MEGAshortscript T7 Transcription kit (Thermo Fisher). *sgRNAs* were mixed with Cas9 Nuclease (Integrated DNA Technologies) to a final concentration of 50-100 ng/µL each *sgRNA* and 1 µg/µL of Cas9 protein, and injected into one-celled zygotes at the volume of 1-2 nL. Progeny of injected F0 generation were screened for the presence of inherited indels resulting in frameshifts or truncations, and these F1 progenies were used to establish stable mutant lines. All *sgRNA* sequences and genotyping primers are listed in Table S1.

#### Immunohistochemistry

##### Drosophila staining

Dissection and immunostaining of adult fly brains were performed according to the FlyLight Protocols (https://www.janelia.org/project-team/flylight/protocols). Adult fly brains were dissected in cold 1xPBS (phosphate buffered saline, Invitrogen), and fixed in 2% paraformaldehyde (PFA, Electron Microscopy Sciences) in 1xPBS for 55 minutes at room temperature. Fixed brains were washed four times with PBSTr (5% Triton X-100 in 1xPBS) while nutating for 20-25 minutes per wash at room temperature. Blocking was performed in 5% normal goat serum (Jackson ImmunoResearch) in PBSTr (0.5% Triton X-100 in 1xPBS) for 1.5-2 hours at room temperature on a nutator. Primary and secondary antibodies were diluted in PBSTr (0.5% Triton X-100) and incubated with samples at 4°C for 48-72 hours. Washes after the primary and secondary antibody incubations were carried out at room temperature with PBSTr for 4 x 25 minutes. Stained samples were mounted with VECASHIELD antifade mounting medium (Vector Laboratories) and stored at 4°C until imaging. Larval CNS dissections were performed in 1xPBS, and fixed in 4% FA/1xPBS for 25 minutes at room temperature. Larval CNS dissections were washed and blocked in PBST-BSA (0.2%Trition X-100 in 1XPBS, with 1% Bovine Serum Albumin). Primary antibodies were diluted in PBST without any blocking agent and were incubated at 4°C for about 48h. Subsequently the samples were washed with PBST 5x for a total time of 1h. Secondary antibodies were diluted in PBST-BSA and incubated at 4°C for about 48 hours, washed with PBST 5x for a total time of 1 hour and mounted in CFM-1 with antifade (Citifluor) and stared at 4°C until analysis on a confocal microscope. Primary antibodies used as follows: rabbit anti-Gat (1:2500, Stork *et al*., 2014), chicken anti-GFP (1:1000, ab13970, abcam), rat anti-mCherry (1:1000, M11217, Thermo Fisher), rabbit anti-RFP (1:1000, 600-401-379, Rockland Inc.), mouse anti-nc82 (1:25, Developmental Studies Hybridoma Bank), anti-Pros (MR1A, 1:150, Developmental Studies Hybridoma Bank). Secondary antibodies used include: Donkey secondary antibodies conjugated to Alexa Fluor 488/647 or Rhodamine Red-X (1:250, Jackson ImmunoResearch), horse anti-mouse DyLight 649 (1:250, DI-2649, Vector Laboratories), and goat anti-rabbit DyLight 649 (1:250, DI-1649, Vector Laboratories).

##### Zebrafish staining

Zebrafish larvae were fixed in 4% PFA/1xPBS at 4°C overnight and then incubated with 150 mM Tris-HCl, pH 9.0 at 70°C for 15 minutes for antigen retrieval (Inoue and Wittbrodt, 2011). Fixed larvae were subsequently permeabilized with 100% acetone at -20°C for 20 minutes, and mouse anti-GS (1:1 prediluted, Sigma-Aldrich) antibody staining was performed as described previously (Chen *et al*., 2020). Stained samples were mounted in VECASHIELD antifade mounting medium (Vector Laboratories) for imaging.

### Whole-mount *in situ* hybridization in zebrafish

To examine the expression patterns of *s1pr1* in zebrafish, we PCR amplified the coding region of *s1pr1* with a T7 promoter sequence in the reverse primer from 2-3 dpf larval stage wild-type cDNA. The anti-sense RNA probes of *s1pr1* were synthesized using Digoxigenin RNA Labeling Kit (Sigma-Aldrich), and whole-mount *in situ* hybridization was performed as previously described (Cunningham and Monk, 2018). 3 dpf zebrafish were fixed in 4% PFA/1xPBS at 4°C overnight and dehydrated with 100% methanol at -20°C for at least 48 hours prior to the following steps. Dehydrated samples were rehydrated through a series of methanol/PBS washes for 10 minutes each at room temperature with an additional 3 x 10 minutes PBSTw (0.1% Tween-20) and treated with proteinase K (1:1000, Bioline) for 20 minutes to increase permeabilization. Samples were then postfixed with 4% PFA/1xPBS for 20 minutes at room temperature, and rinsed 5 x 5 minutes with PBSTw. Pre-hybridization was performed at 65°C in hybridization buffer (50% formamide, 5x SSC, 50 µg/mL heparin, 500 µg/mL transfer RNA, 100 mM citric acid, and 0.1% Tween-20 in dH2O) for at least 4 hours. After prehybridization, samples were incubated with digoxigenin-labeled *s1pr1* RNA probe in hybridization buffer at 65°C overnight. Anti-digoxigenin-AP antibody was used at 1:2000 and followed by alkaline phosphatase staining assay. Stained samples were kept in 80% glycerol at 4°C until photographed.

### Sparse labeling using FLP-out and MARCM in *Drosophila*

To sparsely label adult fly astrocytes with single-cell clones using the FLP-out system, we collected *hsFLPD5.fco;alrm>nlsLexAfl>Gal4co,13xLexAop2-mCD8-GFP,UAS-mCD8-mCherry* or *hsFLPD5.fco;alrm>nlsLexAfl>Gal4co,UAS-mCD8-mCherry;UAS-Lifeact-GFP.W* pupae with the corresponding genotypes (*UAS-RNAi*, wild-type, or *Tre1* mutant backgrounds) at 0-16 h APF and heat shocked at 37°C for 10 minutes at 32-48 h APF. Under these conditions, the FLP-out clones will have the *FRT*-flanking *nlsLexAfl (>nlsLexAfl>*) cassette excised and begin to express *Gal4/UAS*-driven transgenes in the astrocytes. The heat shocked pupae were then incubated at 25°C until eclosion. Adult brain dissections were performed at 1-3 dpe, and subjected to immunostaining described as above before imaging. Similarly larval FLP-out clones were generated using an *alrm*>QF>Gal4co, *UAS-mCD8-mCherry, ftz-FLP stock.* The activity of *ftz*-FLP leads to early and relatively sparse labeling of Gal4 expressing astrocyte (and to some degree ensheathing glia) clones.

MARCM clones were generated by crossing FRTG13 tub-Gal80 *repo*-Gal80; *alrm*-Gal4, UAS-CD8-Cherry, *gcm*-FLP to FRTG13 controls or FRT42B *wun^49^ wun2^Ex34^* or FRT42B *wun^49^ wun2^Ex34^* combined with UAS-*wun* or UAS-*wun2*-GFP on the third chromosome. Note that FRT42B and FRTG13 are the same FRT insertion but are named differently in different research contexts. This *gcm*-FLP MARCM system generates routinely 80-90% of L3 larvae with VNC clones.

### Zebrafish microinjections and mosaic labeling

To label zebrafish astrocytes at the single-cell level, we injected 20-30 pg of *slc1a3b:myrGFP-P2A-H2AmCherry* DNA constructs into one-celled zygotes and allowed the embryos to develop at 28.5°C until desired developmental stages for imaging. We assayed individual spinal cord astrocyte volumes at 6 dpf, and we chose to live-image the dynamics of astrocyte elaboration at 3 dpf when astrocytes are actively growing in the spinal cord (Chen *et al*., 2020).

### Image acquisition and processing

Imaging of fixed fly CNS or zebrafish larvae was performed using Zeiss LSM 880 and LSM 980 with Airyscan 2 confocal microscopes, or an Innovative Imaging Innovations (3i) spinning-disk confocal microscope. For high-resolution astrocyte imaging, confocal z stacks were acquired using the optimal z-interval and around 0.05-0.09 µm/pixel resolution with either a 40x/1.2 Plan-Apochromat Imm Corr or a 40x/1.3 Plan-Apochromat oil objective.

Images were Airyscan processed, images tiles stiched when necessary, and converted into IMARIS format for 3D analysis.

### Zebrafish *in vivo* imaging

To live-image astrocyte morphogenesis in zebrafish, 3 dpf larvae were anesthetized with 0.16 mg/mL of Tricaine in embryo medium and mounted in 1.0% low-melting agarose laterally on a cover slip with extra embryo medium containing Tricaine sealed inside a vacuum grease ring to prevent evaporation. Time-lapse imaging on *slc1a3b:myrGFP-P2A-H2AmCherry*+ astrocyte clones were performed in 40-50 µm z-stack with 0.5 µm z-interval every 10 minutes for 1.5-2 hours. Owing to the z-stack sometimes being insufficient to include an entire astrocyte clone, the z-stack was adjusted to focus on the astrocyte cellular processes where the dynamic astrocytic outgrowths were occurring. Acquired time-lapse images were Airyscan processed, and followed by Bleach correction and 3D drift correction using Fiji (ImageJ) software before analysis.

### Chemical treatments

FTY720 and Ex26 (Tocris Bioscience) were dissolved in DMSO to a stock concentration of 100 mM. To treat zebrafish, the stock solution was diluted to a final concentration of 1 µM with embryo medium with 0.1% DMSO. The *Tg(slc1a3b:myrGFP-P2A-H2AmCherry)* fish were incubated with the diluted solution starting at 2 dpf or 4 dpf for 48 hours, rinsed several times with embryo medium, and incubated with fresh embryo medium until imaging. Control fish were treated with 0.1% DMSO in embryo medium for same period of time. 6 dpf larvae were then mounted dorsally on a cover slip in 1.2% low-melting agarose for imaging.

### Quantification and statistical analysis

To quantify individual astrocyte volumes and densities, we used the Surface module in IMARIS 9 (Bitplane). For astrocyte cellular structure analyses, we used the Filament module in IMARIS 9 with default settings in the software. To quantify the extension and retraction speed of zebrafish astrocyte individual processes, we manually curated the tracking results from frame to frame, and averaged the mean speed for each individually-tracked objects. Quantification of drug treatment experiments on *Tg(slc1a3b:myrGFP-P2A-H2AmCherry)* zebrafish astrocytes were performed by averaging the mean GFP intensity from 4 independent 10 µm x 10 µm regions in the lateral astrocyte processes infiltrated areas in the spinal cord, and similar z-plane images were taken from individual fish. All statistical analyses were carried out using GraphPad Prism 8/9; statistical details, p values, and numbers of analyzed samples are indicated in the figure legends.

## SUPPLEMENTAL INFORMATION

**Figure S1.**
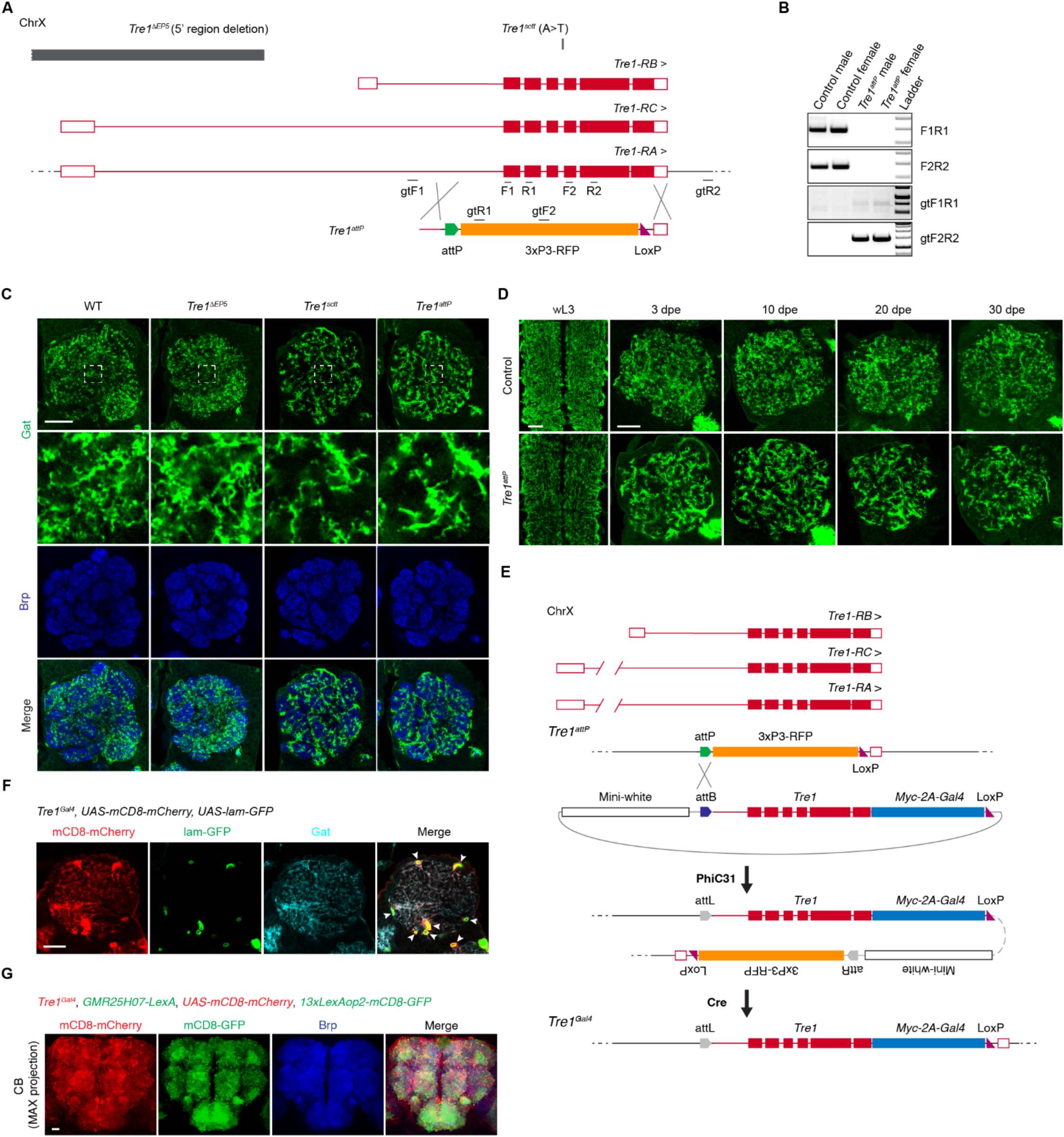
*Tre1* is required for astrocyte morphogenesis in *Drosophila*, related to Figure 1. (A) Schematic of different *Tre1* transcripts and the various *Tre1* genomic mutations. (B) PCR results of the related *Tre1* genotyping primers to validate the *Tre1^attP^* allele in control siblings and in *Tre1^attP^* mutants. (C) Images of astrocytes labeled by α-Gat antibody (green) and the neuropil by α-Brp (blue) in the adult brain ALs. Insets show enlarged views to highlight the astrocyte membrane infiltration differences observed in the corresponding genotypes. N, number of animals. WT, N=9; *Tre1*^D^*^EP5^*, N=7; *Tre1^sctt^*, N=7; and *Tre1^attP^*, N=12. Scale bar, 20 µm. (D) *alrm-Gal4 UAS-mCD8-GFP*-labeled astrocyte membrane in controls and *Tre1^attP^* mutants at wL3 and different adult stages. Representatives of N=4-10 animals independently. Scale bars, 20 µm. (E) Schematic of site-specific integration at the *Tre1^attP^* locus to knock-in the *Tre1* genomic region, a self-cleaving *T2A* peptide sequence, a *Gal4* sequence, and a *LoxP* site. The flanking *LoxP* sites spanning the 3xP3-RFP and mini-white cassette were subsequently eliminated by a Cre stock. (F) Endogenous *Tre1^Gal4^* line drives the expression of *UAS-mCD8-mCherry* and *UAS-lam-GFP* to label cell membrane (red) and nuclei (green) with the astrocyte membrane labeled with α-Gat antibody (cyan) in the ALs (N=7). Arrowheads denote lam-GFP-labeled nuclei in *Tre1* expressing cells colocalized with astrocyte Gat. Scale bar, 20 µm. (G) MAX projection images of *Tre1^Gal4^ UAS-mCD8-mCherry*-expressing cells (red) in comparison with *GMR25H07-LexA 13xLexAop2-mCD8-GFP*-expressing astrocytes (green) in the central brain, also co-labeled with α-Brp (blue) for the neuropil. Scale bar, 20 µm.

**Figure S2.**
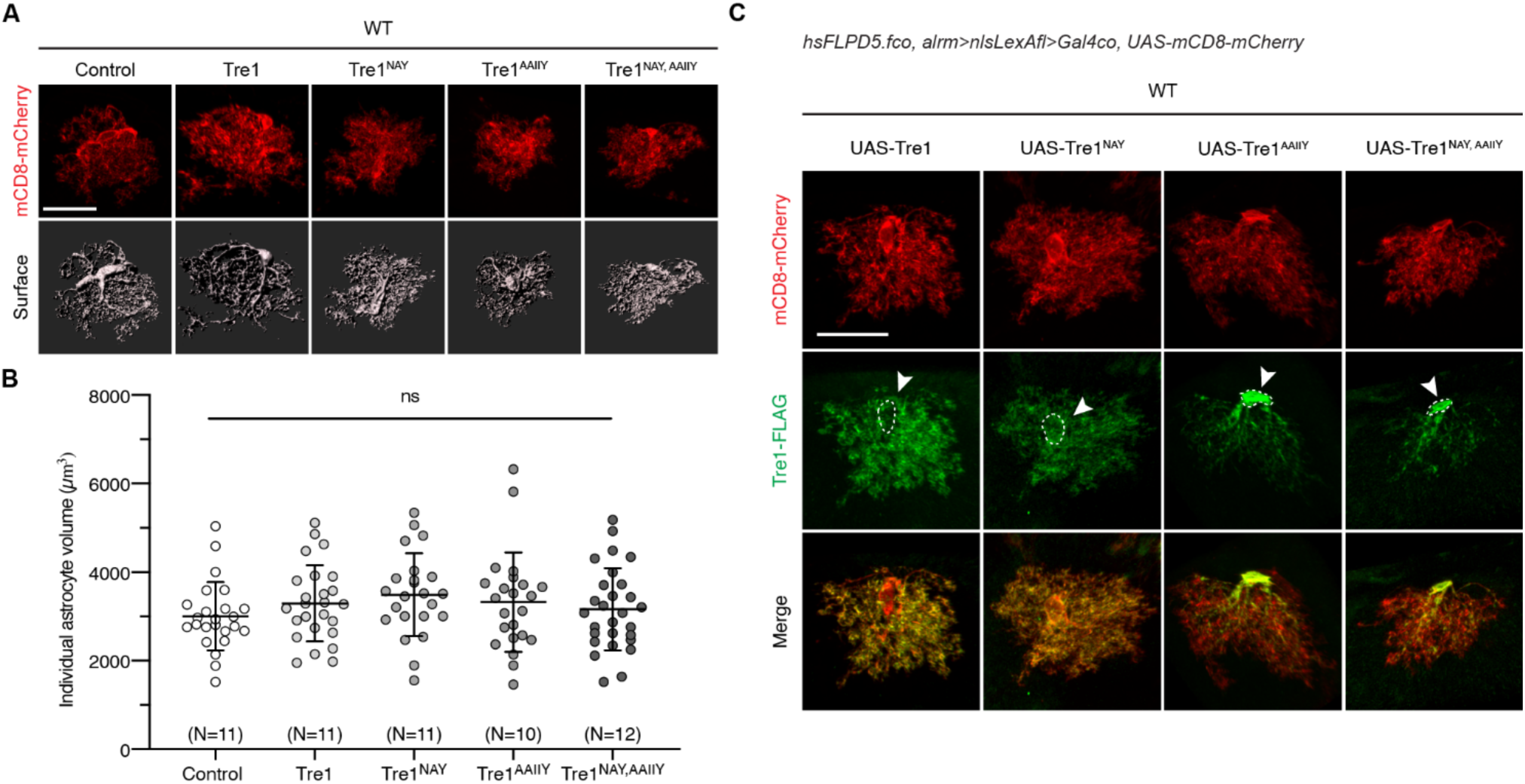
*Tre1* transgenic construct expression in WT astrocytes, related to Figure 2. (A and B) Images (A) of single astrocytes labeled by mCD8-mCherry (red) and the representative IMARIS 3D-rendering surface that expressing different *Tre1* constructs in a WT background, and the quantification (B) of individual astrocyte volumes. Data points represent single astrocytes. N, number of animals. Control, N=11; *Tre1*, N=11; *Tre1^NAY^*, N=11; *Tre1^AAIIY^*, N=10; and *Tre1^NAY,AAIIY^*, N=12. ns, not significant; one-way ANOVA with multiple comparisons. Error bars, mean values ± S.D. Scale bar, 20 µm. (C) Images of individual astrocyte clones labeled by mCD8-mCherry (red) expressing different *Tre1* constructs in a WT background also co-stained with a-FLAG antibody (green) to localize Tre1 protein subcellular distribution. Dashed circles and arrowheads mark the astrocyte cell body regions. Scale bar, 20 µm.

**Figure S3.**
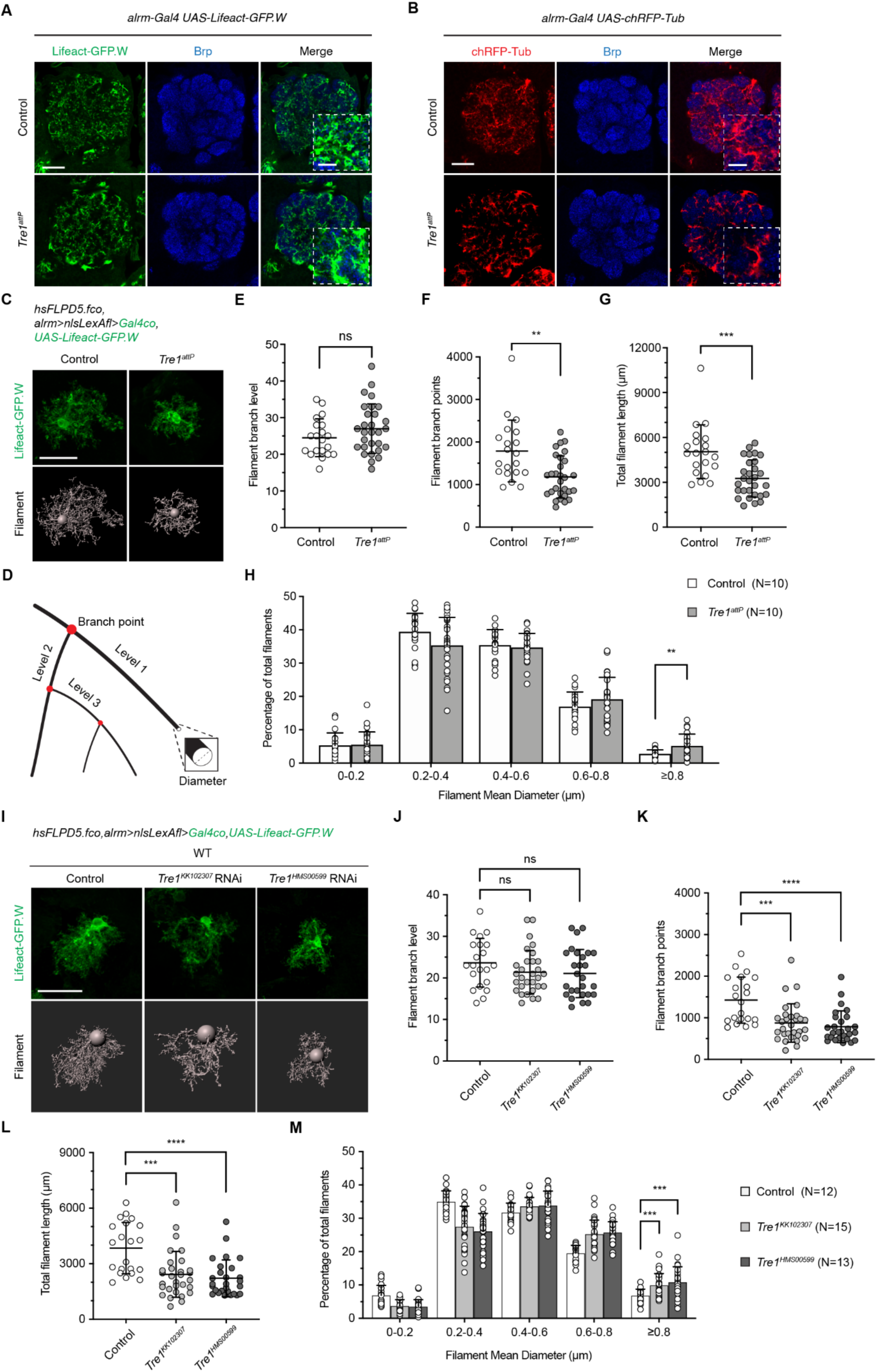
Loss of *Tre1* leads to cytoskeleton defects in astrocytes, related to Figure 4. (A) Images of AL astrocyte actin cytoskeleton structure labeled by *alrm-Gal4 UAS-Lifeact-GFP.W* (green) with Brp-labeled neuropil (blue) in control (N=8) and *Tre1^attP^* mutants (N=10). N, number of animals. Insets show enlarged views of the merged images. Scale bars, 20 µm and 5 µm (insets). (B) Images of AL astrocyte microtubule structure labeled by *alrm-Gal4 UAS-chRFP-Tub* (red) with α-Brp (blue) stained neuropil in control (N=8) and *Tre1^attP^* mutants (N=5). N, number of animals. Insets show enlarged views of the merge images. Scale bars, 20 µm and 5 µm (insets). (C) Images of single-cell astrocytes labeled with actin marker Lifeact-GFP.W (green) using the FLP-out system and the representative IMARIS 3D-rendering filament structures (grey) in control and *Tre1^attP^* mutants. Scale bar, 20 µm. (D) Schematic of IMARIS/filament-detected cytoskeleton branch level, branch point, and mean diameter. (E-H) Quantification of single-cell astrocyte cytoskeletal branch level (E), branch points (F), total length (G), and distribution of mean diameter (H) in control (N=10) and *Tre1^attP^* mutants (N=10). N, number of animals. Data points represent single astrocytes. ns, not significant; **, p<0.01; ***, p<0.001; unpaired t test. Error bars, mean values ± S.D. (I) Images of single astrocytes labeled with actin marker Lifeact-GFP.W (green) and the representative IMARIS 3D-rendering filament structures (grey) in control, *Tre1^KK102307^ RNAi*, and *Tre1^HMS00599^ RNAi*. Scale bar, 20 µm. (J-M) Quantification of single-cell astrocyte cytoskeletal branch level (J), branch points (K), total length (L), and distribution of mean diameter (M) in control (N=12), *Tre1^KK102307^ RNAi* (N=15), and *Tre1^HMS00599^ RNAi* (N=13). N, number of animals. Data points represent single astrocytes. ns, not significant; ***, p<0.001; ****, p<0.0001; one-way ANOVA with multiple comparisons. Error bars, mean values ± S.D.

**Figure S4.**
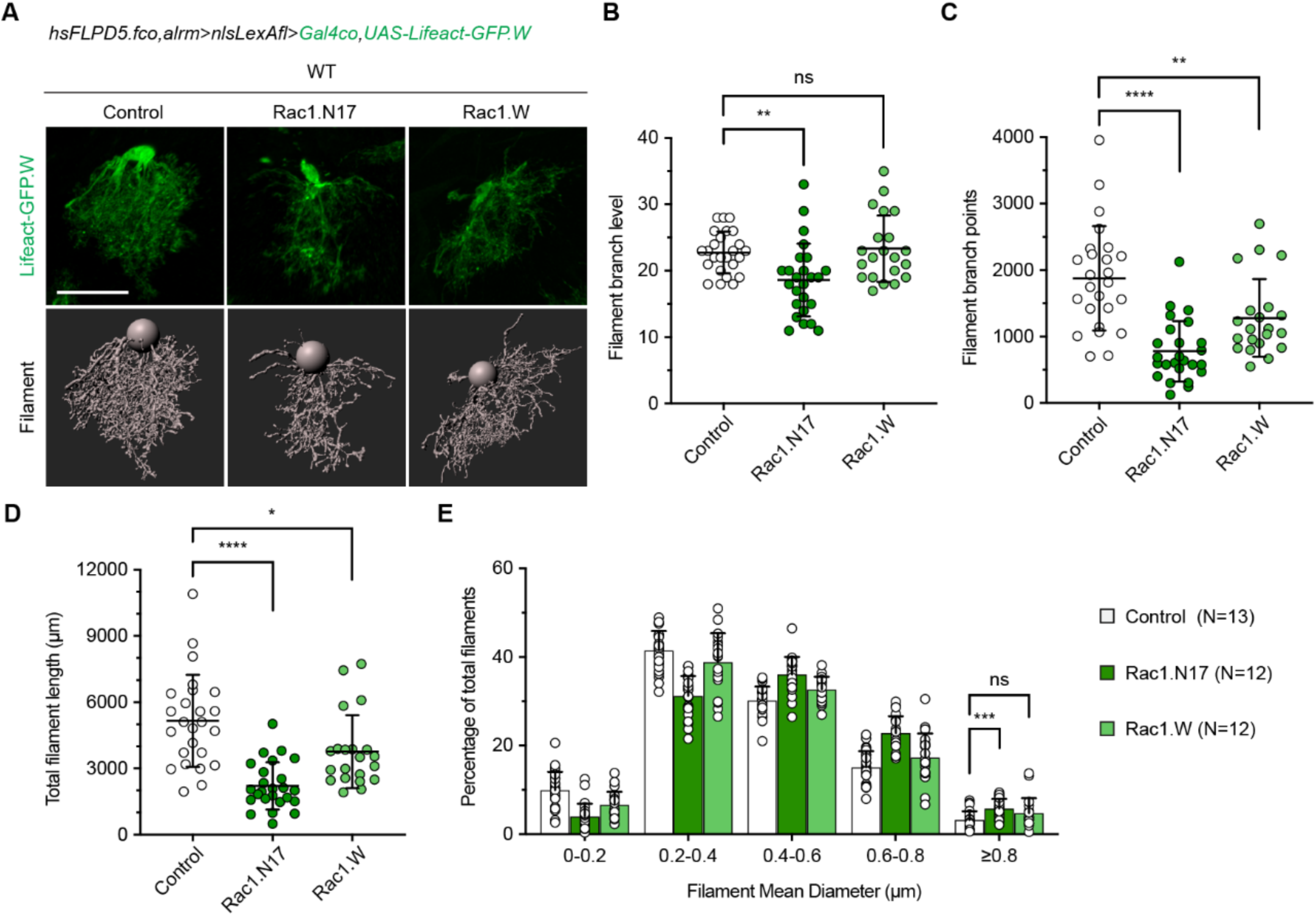
Proper Rac1 activity in astrocytes is crucial for normal astrocyte development, related to Figure 4. (A) Images of single-cell astrocytes labeled with Lifeact-GFP.W (green) and the representative IMARIS 3D-rendering filament structures (grey) in WT backgrounds that express *Rac1.N17* or *Rac1.W*. Scale bar, 20 µm. (B-E) Quantification of single-cell astrocyte cytoskeletal branch level (B), branch points (C), total length (D), and distribution of mean diameter (E) in control (N=13), *Rac1.N17* (N=12), and *Rac1.W* (N=12). N, number of animals. Data points represent single astrocytes. ns, not significant; *, p<0.05; **, p<0.01; ***, p<0.001; ****, p<0.0001; one-way ANOVA with multiple comparisons. Error bars, mean values ± S.D.

**Figure S5.**
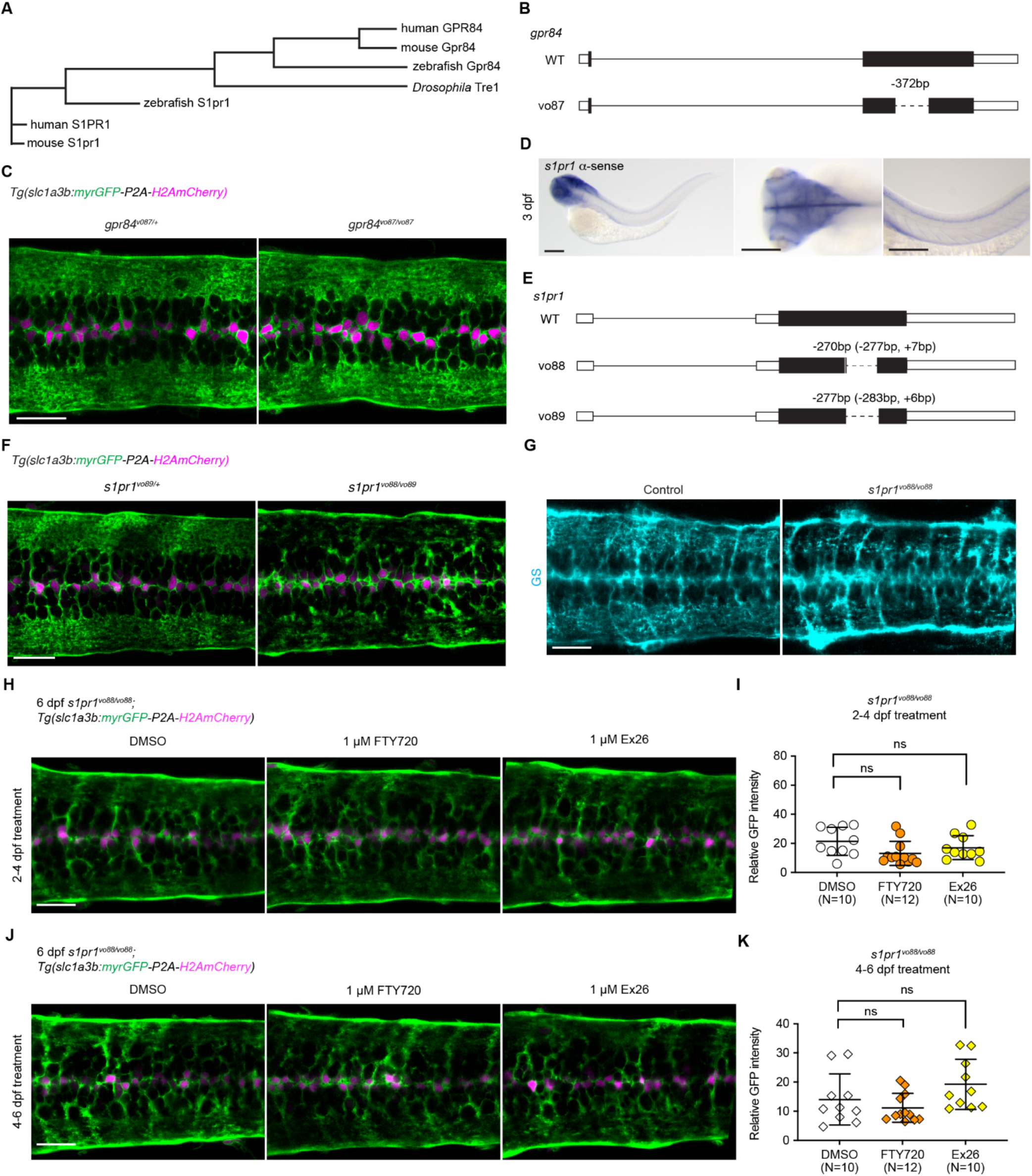
*s1pr1*, but not *gpr84*, is required for zebrafish astrocyte morphogenesis, related to Figure 5. (A) Phylogenetic comparison of *Drosophila* Tre1 protein with the most closely related GPCR Gpr84 and the same class A GPCR S1pr1 in zebrafish, mouse, and human. (B) Schematic of the *gpr84^vo87^* mutant allele generated by two sgRNAs-mediated CRISPR/Cas9 approaches in zebrafish. The *vo87* mutation results in a deletion of 372 bp in the coding region of the *gpr84* gene. (C) Images of 6 dpf *gpr84^vo87/+^* heterozygous controls (N=4) and *gpr84^vo87/vo87^* homozygous mutants (N=4) spinal cord astrocytes labeled with myrGFP (green) and H2AmCherry (magenta) using the *Tg(slc1a3b:myrGFP-P2A-H2AmCherry*) transgene. Scale bar, 20 µm. (D) Whole mount *in situ* hybridization results of 3 dpf zebrafish larvae using an *s1pr1* a-sense probe. Representatives of N≥30 larvae analyzed. Scale bars, 200 µm. (E) Schematics of the *s1pr1^vo88^* and *s1pr1^vo89^* mutant alleles generated by two sgRNAs-mediated CRISPR/Cas9 approaches in zebrafish. The *vo88* mutation results in 270 bp (-277,+7) deletion, and *vo89* results in 277 bp (-283, +6) deletion in the coding region of *s1pr1* gene. These two alleles were generated from independent CRISPR/Cas9 injection experiments. (F) Images of spinal cord astrocytes in 6 dpf *Tg(slc1a3b:myrGFP-P2A-H2AmCherry*) transgenic *s1pr1^vo89/+^* heterozygous controls (N=12) and *s1pr1^vo88/vo89^* trans-heterozygous mutants (N=7). Scale bar, 20 µm. (G) Images of 6 dpf control (N=11) and *s1pr1^vo88/vo88^* mutant (N=11) spinal cord stained with astrocyte marker a-GS antibody (cyan). Scale bar, 20 µm. (H and J) Images of 6 dpf *s1pr1^vo88/vo88^* mutant spinal cord astrocytes after treated with DMSO, 1 µM FTY720, or 1 µM Ex26 at 2-4 dpf (H) or 4-6 dpf (J) in the *Tg(slc1a3b:myrGFP-P2A-H2AmCherry)* transgenic background. Scale bar, 20 µm. (I) Quantification of relative GFP intensity at 6 dpf *s1pr1^vo88/vo88^* mutants in the astrocyte process-enriched regions in DMSO (N=10), FTY720 (N=12), and Ex26 (N=10) after 2-4 dpf treatment. N, number of animals. Data points represent single fish. ns, not significant; one-way ANOVA with multiple comparisons. (K) Quantification of relative GFP intensity at 6 dpf *s1pr1^vo88/vo88^* mutants in the astrocyte process-enriched regions in DMSO (N=10), FTY720 (N=12), and Ex26 (N=10) after 4-6 dpf treatment. N, number of animals. Data points represent single fish. ns, not significant; one-way ANOVA with multiple comparisons.

**Figure S6.**
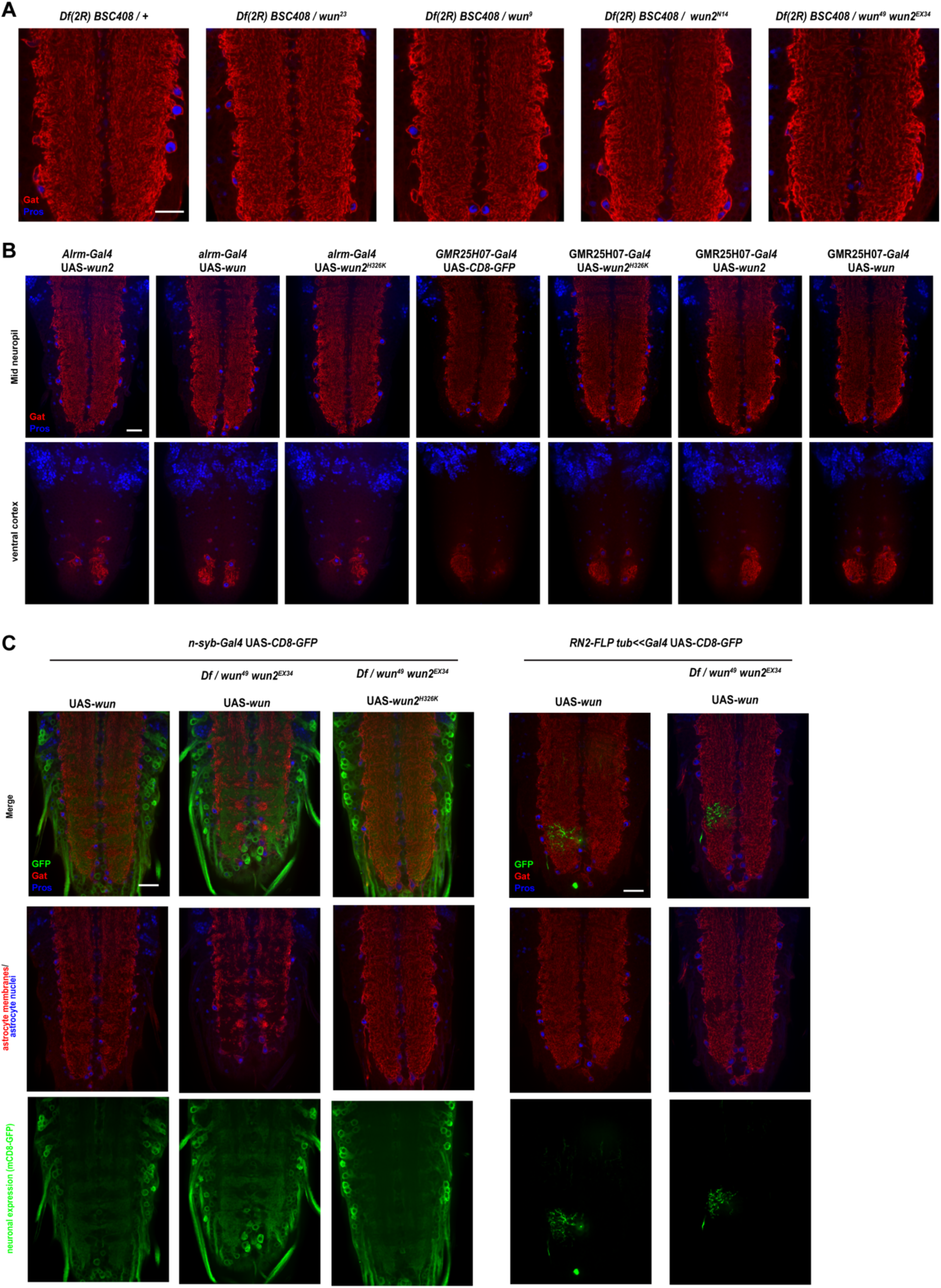
*wun/wun2* mutant and misexpression phenotypes, related to Figure 6. (A) Confocal single section images of *Df(2R) BSC408/+* control (N= 6), *Df(2R) BSC408/wun^23^* (N= 7), *Df(2R) BSC408/wun^9^* (N= 6), *Df(2R) BSC408/wun2^N14^* (N= 9), and *Df(2R) BSC408/wun^49^,wun^EX34^* (N= 8) VNC astrocytes with membrane labeled by α-Gat antibody (red) and nuclei labeled by α-Pros antibody (blue) in L3. Scale bar, 20 µm. (B) Images of VNC astrocytes labeled by Gat (red) and Pros (blue) that express different *wunen* transgene constructs using either *alrm-Gal4* or the stronger astrocyte driver *GMR25H07-Gal4*. N=3-4 animals. Scale bar, 20 µm. (C) Single section confocal images of L3 stage VNC with astrocyte membranes labeled by α-Gat (red) and nuclei labeled by α-Pros (blue). UAS-*wun* or UAS-*wun2^H326K^* are overexpressed in wild-type or *Df(2R)w45-19g*/*wun^49^*,*wun2^EX34^* backgrounds either pan-neuronally with *n-syb-*Gal4or in single cell neuronal clones using *RN2-FLP tub<<Gal4* labeled with UAS-mCD8-GFP (green). *n-syb*-Gal4 UAS-CD8-GFP UAS-*wun* (N=3); *Df(2R)w45-19g/wun^49^,wun^EX34^*; *n-syb*-Gal4 UAS-CD8-GFP UAS-*wun* (N=8); *Df(2R)w45-19g/wun^49^,wun^EX34^*; *n-syb*-Gal4 UAS-CD8-GFP UAS-*wun2^H326K^* (N= 6); *RN2-FLP tub<<Gal4* UAS-CD8-GFP UAS-wun (N=6); *Df(2R)w45-19g/wun^49^,wun^EX34^*; *RN2-FLP tub<<Gal4* UAS-CD8-GFP UAS-wun (N=10); Scale bars, 20 µm.

**Figure S7.**
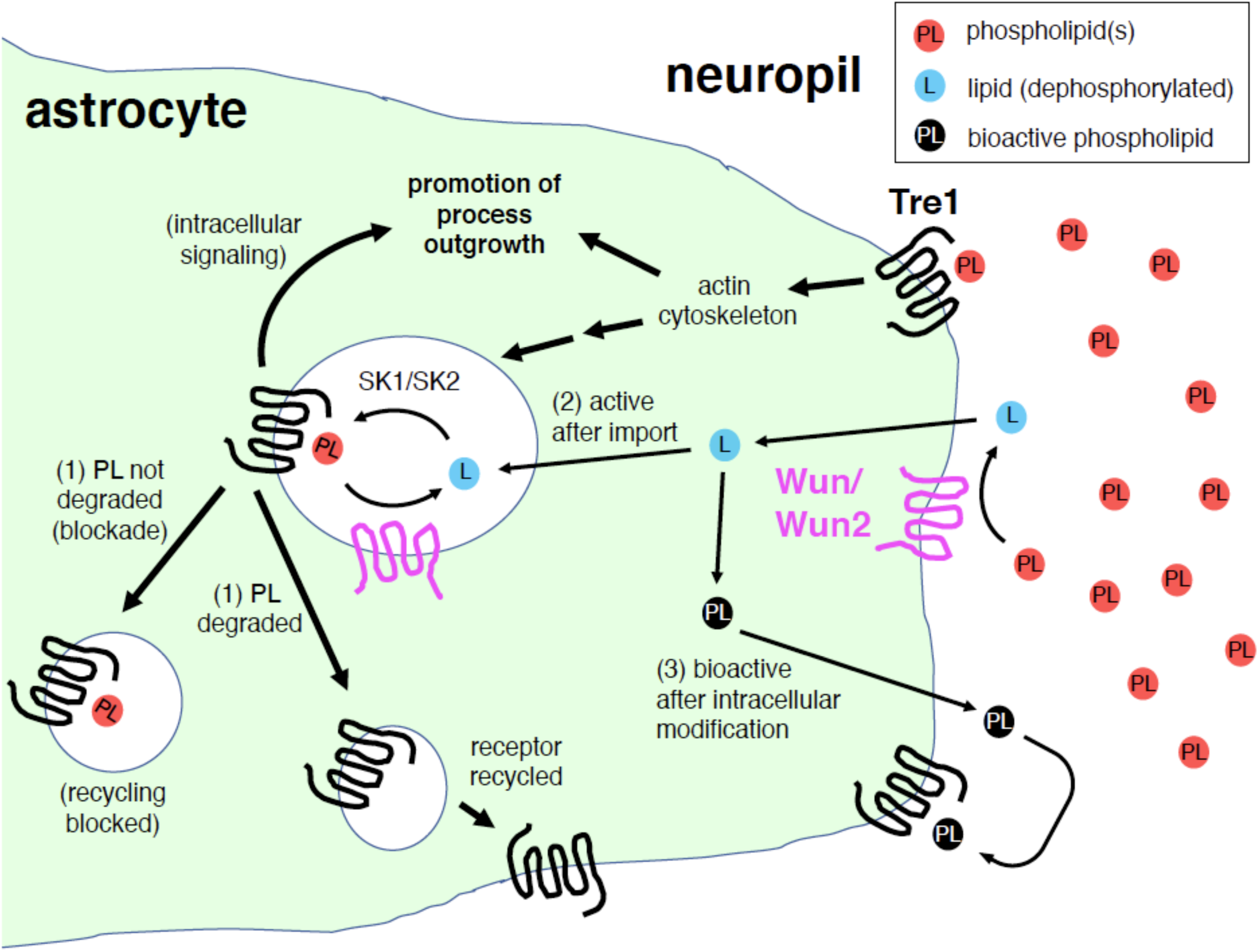
Model for Tre1 and Wun/Wun2 signaling. Astrocyte growth-promoting phospholipids (PL) are distributed throughout the neuropil, potentially secreted by neurons. PL binding to Tre1 promotes process outgrowth through regulation of actin cytoskeletal dynamics, which also leads to Tre1 receptor internalization. We envision three models (not mutually exclusive) for signaling: (1) Signaling is maintained by Wun/Wun2-dependent degradation of PL bound to Tre1 by enabling efficient Tre1 receptor recycling to the plasma membrane after PL degradation. (2) PL is dephosphorylated by Wun/Wun2, thereby allowing entry into the cell, where it is re-phosphorylated by a lipid kinase (e.g. SK1/SK2) to activate Tre1 and drive intracellular signaling. (3) PL is dephosphorylated by Wun/Wun2, enters the cell, where it is rephosphorylated into its bioactive form, which is then transported outside the cell to activate Tre1 receptors locally. Overexpression of Wun/Wun2 in neurons is able to suppress ingrowth of astrocyte processes by depleting local Pls. Overexpression of Wun or Wun2 in a single cell clone in a *wun, wun2* double mutant is able to drive massive overgrowth because only the Wun-expressing astrocyte is able to process growth­ promoting Pls.

**Video 51.** 3D-rendering of single astrocyte actin cytoskeleton labeled by Lifeact-GFP.W (green) and the !MARlS-generated filament structure in WT control and *Tre1affP* mutants. Scale bar, 71-Jm. Related to Figure 4.

**Video 52.** Time-lapse movie of single-cell astrocyte membrane dynamics labeled by slc1a3b:myrGFP at 3 dpf

**Table S1.**
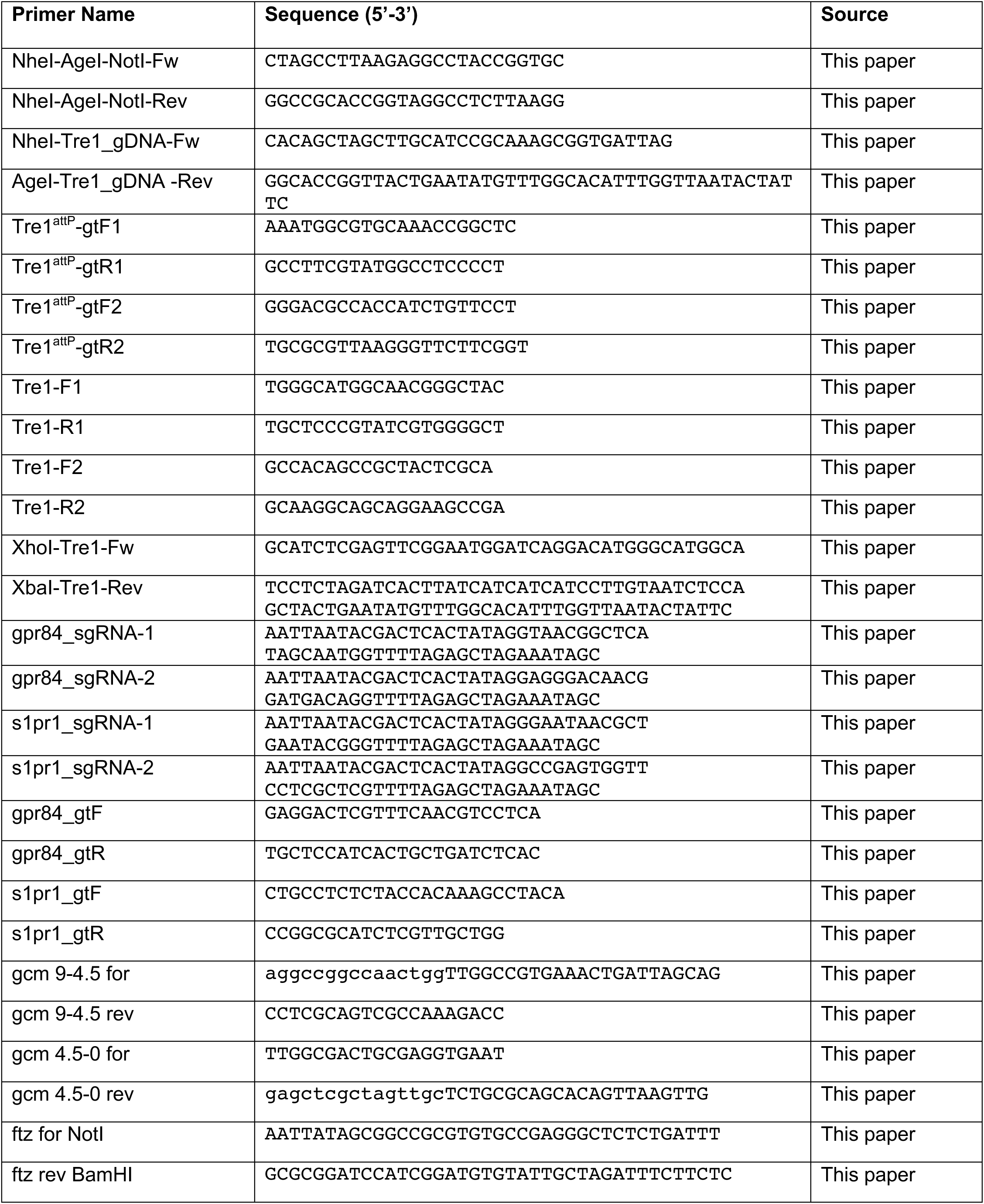
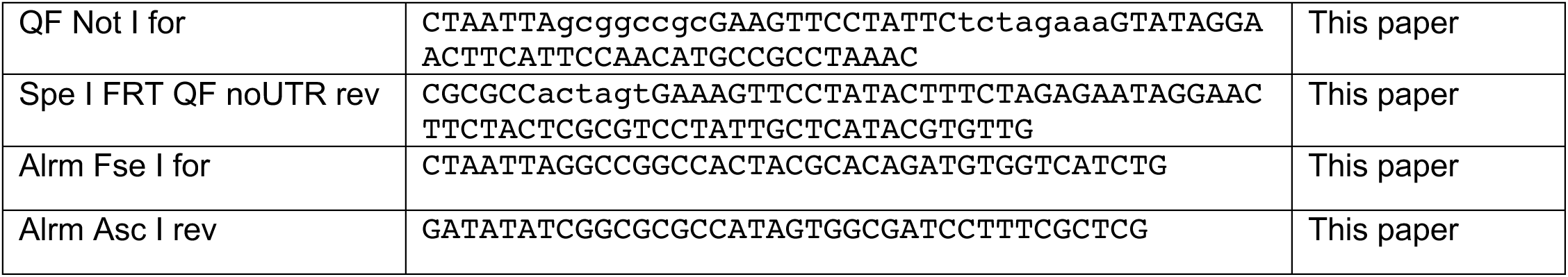
List of primer sequence used in this study, related to Methods section.

